# A Single Genomic Region Controls Primocane Fruiting in Tetraploid Blackberry

**DOI:** 10.64898/2025.12.15.694498

**Authors:** Alexander Silva, Isabella Vaughn, T. Mason Chizk, Lacy Nelson, Carmen Johns, Ellen Thompson, Nahla Bassil, Michael Hardigan, John Clark, Tomas Bruna, Marcelo Mollinari, Margaret Worthington

## Abstract

The fresh-market blackberry (*Rubus* subgenus *Rubus*) industry has expanded dramatically in the past two decades, driven in part by improved cultivars. Introgression of the primocane-fruiting (PF; annual flowering) trait into elite germplasm has enabled dual cropping in a single year, season extension, and cultivation in tropical and subtropical regions. Despite its economic performance, the genetic basis of PF is not well understood. It has been proposed that the PF trait is controlled by a major recessive locus, but its genomic location is unclear. Here, a genome-wide association study (GWAS) of 365 tetraploid blackberry genotypes identified a single genomic region on chromosome Ra03 (∼33 Mb) strongly associated with PF. Genetic linkage analysis in a biparental population confirmed that the same interval (32-35 Mb) was linked to the PF phenotype. Ten putative candidate genes were identified in this region. Allele mining using whole-genome resequencing of 17 genotypes highlighted two high-priority candidates: a CCCH-type zinc finger gene and a ubiquitin-specific protease gene. Use of an improved *Rubus argutus* ‘Hillquist’ genome annotation (v1.2) enabled refined variant interpretation, including identification of regulatory 3′ UTR polymorphisms in the zinc finger homolog. Two diagnostic KASP markers (*PF1* and *PF2*), designed from the most significant GWAS SNPs, predicted the PF phenotype with over 96% accuracy in a validation panel of 494 tetraploid blackberries from multiple breeding programs. Together, these results provide the first high-resolution mapping of the PF locus in blackberry, identify candidate genes for flowering regulation in *Rubus*, and deliver diagnostic markers that can be immediately deployed in breeding programs.

## Introduction

Blackberry (*Rubus* subgenus *Rubus*) is a specialty crop of increasing economic significance due to rising consumption, expanded marketing initiatives, and advancements in cultivar development (Clark and Finn, 2014). Blackberries are herbaceous or semi-woody plants with a perennial crown root system and biennial canes. The first-year canes (primocanes), are typically vegetative, while the second-year canes (floricanes), produce flowers and fruits after a period of dormancy requiring varying degrees of chilling hours (often during winter in temperate environments). Genotypes that produce fruit exclusively on floricanes are known as floricane-fruiting (FF) or biennial-flowering cultivars. In contrast, primocane-fruiting (PF) or annual-flowering cultivars flower and fruit on new canes produced each year (Clark, 2008) (Fig. 1).

**Figure 1.**
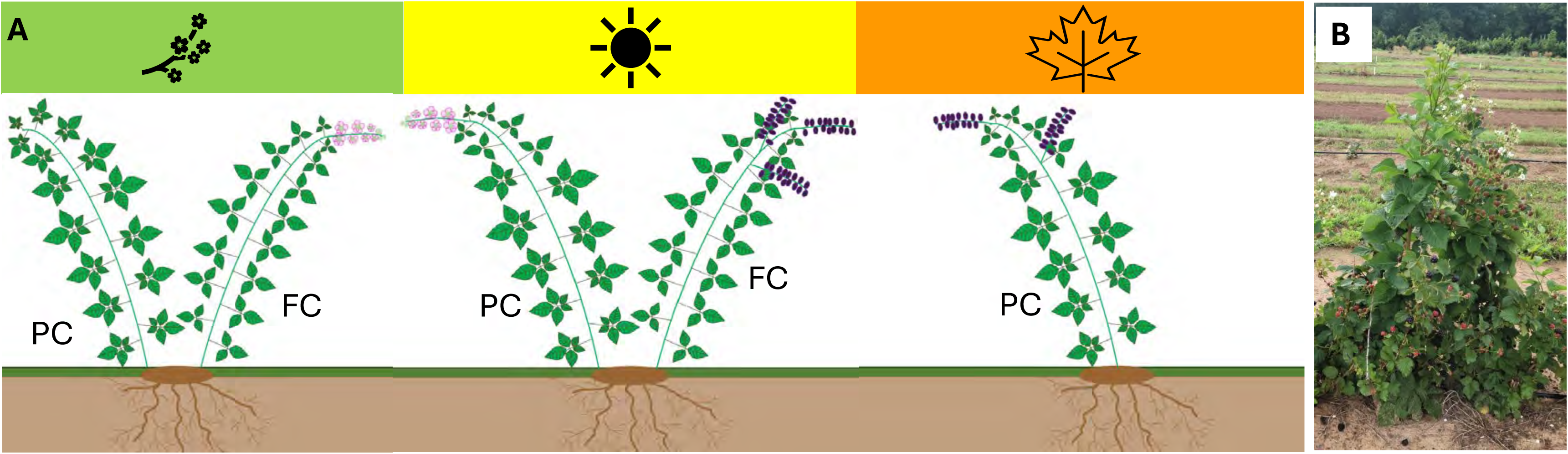
Diagram of flowering pattern in primocane-fruiting genotypes (A). While primocanes (PC) are vegetative in Floricane-Fruiting (FF) blackberry cultivars, they flower during summer in Primocane-Fruiting (PF) cultivars and produce fruit in late summer and fall. Flowering in floricanes (FC) occurs during spring after a period of dormancy, with fruit harvest in summer. Therefore, PF cultivars allow two harvest seasons, one in summer and one in fall. (B) PF genotype ‘S442’ showing unripe fruit and flowers on PCs at the top with ripe fruit on FC below.

Primocane-fruiting is a trait also present in other *Rubus* crops and has been critical for the expansion of red raspberry (*R. ideaus*) production (Clark et al. 2007). In blackberry, the PF trait was initially found in a wild *R. argutus* accession named ‘Hillquist’ (PI 553951). This diploid blackberry accession was used as the initial donor parent to develop tetraploid PF cultivars at the Fruit Breeding Program of the University of Arkansas System Division of Agriculture (UADA), which released the first PF cultivars (Prime-Jan^®^ and Prime-Jim^®^) in 2004 (Clark et al., 2005).

PF cultivars have some advantages over traditional floricane-fruiting cultivars, such as extended production season in the autumn or a double-cropping system (a first harvest season on floricanes in early summer and a second season on primocanes in late summer and fall), scheduled production based on primocane management, avoidance of winter injury in colder areas, and expanded production in regions where chilling requirements are not entirely fulfilled (Clark and Finn, 2014).

Although PF blackberry cultivars are widely adopted in the fresh-market raspberry and blackberry industries, the molecular mechanism and genes controlling this trait have not been well elucidated in either species. Several studies have been conducted to understand the inheritance of primocane fruiting in tetraploid blackberries. Lopez-Medina et al. (2000) evaluated PF segregation ratios in a full diallel crossing scheme and proposed that PF is controlled by a single recessive gene. In addition, Castro et al. (2013) constructed the first genetic map of tetraploid blackberry from a cross between Prime-Jim^®^ (PF cultivar) and ‘Arapaho’ (FF cultivar) using 119 simple sequence repeat (SSR) markers. They identified a locus linked with the PF trait in this segregating population on linkage group (LG) 7 between the markers RH_MEa0006aC04-175 and RH_MEa0007aG06-152. However, it was later shown that most of the markers on LG7, including the flanking markers to the PF locus, aligned on chromosome 2 in the black raspberry (*R. occidentalis*) genome (VanBuren et al. 2016) and the blackberry ‘Hillquist’ reference genome (Brůna et al. 2023). The genetic control of the PF trait has been investigated in red raspberry as well; Jibran et al. (2019) reported two quantitative trait loci associated with PF in red raspberry, *RiAF3* and *RiAF4* on linkage groups 3 and 4, respectively, using a biparental population from the cross between the PF accession NC493 (*R. parvifolius* x *R. idaeus* ‘Cherokee’) and the FF *R. idaeus* genotype ‘Chilliwack’. This finding suggested that PF in raspberries is a complex trait controlled by more than one locus, in contrast to previous findings in tetraploid blackberries.

The development of genomic resources for blackberries is critical for the elucidation of the genetic control of important agronomic traits such as PF. Two reference quality assemblies of blackberry have recently become available. The first, from the diploid donor of the PF trait (*R. argutus* cv. ‘Hillquist’) was assembled at the chromosome scale using Pacific Biosciences long reads, with 38,503 predicted protein-coding genes, 72% of which were functionally annotated (Brůna et al. 2023). The second, from the PF thornless tetraploid blackberry selection BL1, was assembled and haplotype-phased at the chromosome scale using Oxford Nanopore long reads and Hi-C scaffolding, yielded 27 pseudochromosomes and 87,968 predicted protein-coding genes, 82% with functional annotation (Paudel et al. 2025). Together, these genomic resources provide a robust foundation for the identification of loci and candidate genes controlling PF and other complex traits in blackberry.

Next-generation sequencing (NGS) technologies have enabled the generation of large sets of markers with sufficient read depth to accurately estimate allele dosages in polyploid crops using software such as the R package Updog (Gerard et al. 2018), which accounts for variable read depth, sequencing errors, allele bias, and overdispersion. Recently, Capture-Seq genotyping (RAPiD Genomics) was performed using 35,054 biotinylated probes distributed across the ‘Hillquist’ reference genome to identify 124,564 biallelic SNPs across a diverse panel of 502 blackberry genotypes (Chizk et al. 2023). This tetraploid blackberry panel and set of SNPs have been used to identify loci controlling fruit firmness, red drupelet reversion, soluble solid content, pH, acidity, and the prickle-free trait in tetraploid blackberry (Chizk et al. 2023; Johns et al. 2025; Godwin et al. 2025).

Genome-wide association studies (GWAS) and linkage mapping have been extensively used across various species to identify loci associated with diverse traits. The development of the GWASpoly software (Rosyara et al. 2016), designed specifically for autopolyploids to model different types of polyploid gene action, including additive and simplex dominant, has enabled the identification of genomic regions in crops such as blueberry (Ferrão et al. 2018), potato (Sharma et al. 2018), sweet potato (Wilson et al. 2021), sugarcane (Saavedra-Díaz et al. 2024), and blackberry (Chizk et al. 2023; Johns et al. 2025; Godwin et al. 2025). Similarly, the development of the R package MAPpoly (Mollinari and Garcia, 2019) has facilitated the efficient construction of linkage maps using thousands of SNP markers in autopolyploid species, such as in blueberry (Cappai et al. 2020), sweetpotato (Mollinari et al. 2020; Oloka et al. 2021), potato (da Silva et al. 2021), and rose (Lau et al. 2022).

In this study, a panel of 365 diverse tetraploid blackberry genotypes and a segregating biparental population were used to identify genomic regions linked to the PF trait through GWAS and linkage mapping. Based on these analyses, diagnostic Kompetitive Allele-Specific PCR (KASP) markers were developed and validated in a diverse blackberry panel. The most predictive KASP markers have practical applications in breeding programs for marker-assisted selection.

Furthermore, identification of candidate genes and allele mining using whole-genome sequence data and an improved annotation of *R. argutus* cv. ‘Hillquist’ enhances the understanding of the genetic control underlying the PF trait.

## Materials and Methods

### Plant material

A diverse panel of 365 tetraploid blackberry cultivars and selections from the UADA breeding program were classified as PF or FF based on the presence or absence of flowers on first-year canes. Genotypes with at least 20% of primocanes in the plot producing flowers in at least one year between 2019 and 2023 were considered PF genotypes. Each cultivar and selection was maintained in a 6 m plot with an average of 10 plants per plot at the UADA Fruit Research Station (FRS), Clarksville, AR, located at 35°31’5” N and long. 93°24’12” W. Blackberry plots received regular maintenance consisting of pruning, tipping, integrated pest management, and irrigation as recommended in the Southeast Regional Caneberry Production Guide (Fernandez et al. 2023).

As a complementary strategy, genetic linkage analysis was conducted to identify genomic regions associated with the PF trait. A blackberry biparental mapping population was generated in 2019 by crossing the FF selection ‘S16’ and the PF selection ‘S242’. The resulting progeny were planted at 0.6 m spacing at FRS in Spring 2020. Each seedling was screened for the presence or absence of flowers on primocanes during June and July of 2022 and 2023.

### Genotyping and SNP calling

The tetraploid blackberry diversity panel was previously genotyped through the Capture-Seq technology by RAPiD Genomics (Gainesville, FL), as described in Chizk et al. (2023). Briefly, DNA from young leaf tissue of each genotype was extracted using a cetyltrimethylammonium bromide (CTAB) protocol modified from Porebski et al. (1997). A Qubit dsDNA assay kit (Invitrogen, Carlsbad, CA) was used to quantify DNA concentration. A set of 35,054 custom biotinylated 120-mer probes distributed along the ‘Hillquist’ (*R. argutus*) reference genome (Brůna et al. 2023) were used to sequence each genotype. DNA libraries were sequenced using an Illumina HiSeq2000 instrument with an average coverage of 150x. Illumina paired-end reads were cleaned, trimmed, and aligned to the reference genome using MOSAIK v2.2.30 (Lee et al., 2014), and diploid variants were detected using Freebayes v1.3.1 (Garrison and Marth, 2012).

VCFtools (Danecek et al., 2011) was used to select biallelic SNPs with a minor allele frequency ≥ 0.01. Tetraploid allele dosage was estimated using the multidog function from Updog v2.0.3 (Gerard et al. 2018) based on the filtered VCF file.

The biparental population was genotyped using Allegro^®^ targeted genotyping (Tecan Genomics, Redwood City, CA, USA. DNA from young leaf tissue of parents and 165 seedling progenies was extracted as previously described. A custom set of 46,485 probes was developed to target 25,000 sites across the *R. argutus* ‘Hillquist’ reference genome. Of these targets, 23,076 were fully covered and 433 were partially covered. Sequencing libraries were prepared following the Allegro protocol and sequenced using an Illumina NextSeq 2000 to generate 100-bp single-end reads. The sequencing reads were cleaned and trimmed using Trimmomatic v0.39 (Bolger et al. 2014) and aligned to the ‘Hillquist’ reference genome using Burrows-Wheeler Aligner (bwa)-mem2 v0.7.17 software (Li and Durbin, 2009). Genetic variants were called using GATK v4.2.6.1 HaplotypeCaller with the tetraploid option (-ploidy 4), followed by joint genotyping for the whole population with the GenotypeGVCFs tool in GATK (Poplin et al., 2018). SNPs with low confidence were filtered using the following parameters: Phred-scaled quality score below 30 (QUAL < 30), normalized quality score below 2 (QUAL/DP < 2.0), mapping quality below 40 (MQ < 40.0), or depth of coverage less than 50 for each sample (FMT/DP < 50).

### Genome-wide association analysis

GWAS was conducted using the GWASpoly v2.30 (Rosyara et al. 2016) R package under the K model to control population structure. The K matrix was constructed using the leave-one-chromosome-out (LOCO) method, in which a covariance matrix is calculated for each chromosome based on the markers from all other chromosomes. Based on previous research demonstrating that PF is controlled by a single recessive locus in blackberry (Castro et al., 2013; Lopez-Medina et al. 2000), the complete dominance (1-dom) model of gene action was tested. A maximum genotype frequency of 0.98 was set. The “M.eff” method was used to establish - log10(p) significance thresholds of 4.93, and 5.5 for the complete alternative allele dominance and complete reference allele dominance models, respectively, to control the genome-wide false discovery rate. QQ-plots of observed versus expected p-values under the null hypothesis were constructed to assess potential inflation of the test statistics.

### Linkage mapping for the primocane-fruiting locus

A genetic linkage map for the biparental population was constructed using the R package MAPpoly2 (https://github.com/mmollina/mappoly2; Mollinari and Garcia, 2019). SNPs and progenies with more than 10% missing data were excluded. The chi-square (χ2) test was used to assess segregation distortion, assuming only random chromosome bivalent pairing (no double reduction). Redundant markers were automatically removed before computing the pairwise recombination fraction between all the selected markers using a two-point analysis. The recombination fraction between the PF phenotype and each molecular marker was also calculated. Markers were grouped using two sources of information: the chromosome information from the physical map of the *R. argutus* cv. ‘Hillquist’ genome, and the recombination fractions between markers using the Unweighted Pair Group Method with Arithmetic Mean (UPGMA) clustering method. The function “make_sequence” of MAPpoly2 was used to combine the UPGMA-derived groups and the chromosome assignments to ensure the most consistent and supported grouping.

The multidimensional scaling (MDS) method, based on pairwise recombination fractions, was used to determine the order of the markers within each group. The genome-based ordering was included in the analysis to refine the order of markers. Linkage phase was estimated using the function “pairwise_phasing” which utilizes pairwise recombination information to identify a set of linkage phase configurations for each parent. Individual genetic maps were constructed for each parent using a multilocus approach in which the entire chromosome structure is considered; misplaced markers or incorrect linkage phases can be detected in this approach as an inflation of the expected map length. Individual maps were integrated using a Hidden Markov Model (HMM) model, considering a global genotyping error of 5%. MAPpoly2 also calculated homologous pairing probabilities, allowing for detection of preferential pairing during meiosis.

### Structural annotation of *R. argutus* cv. ‘Hillquist’ v.1.2

An updated gene annotation was generated for the *R. argutus* ‘Hillquist’ genome assembly in collaboration with the Joint Genome Institute (JGI). While no new genomic or transcriptomic sequence data were generated, existing Iso-Seq and RNA-Seq datasets (Bruna et al. 2023) were reanalyzed in the pipeline, resulting in substantially improved gene models, including more complete untranslated regions (UTRs), refined exon and intron structures, and updated functional annotations. Transcript assemblies were derived from ∼136M pairs of 2 x 100 stranded paired-end Illumina RNA-seq reads. Reads were aligned to the genome using GSNAP within the PERTRAN framework, which constructs splice alignment graphs after alignment validation, realignment, and correction (Wu and Nacu, 2010). Approximately 2.5M PacBio Iso-Seq non-chimeric CCSs were processed with a genome-guided correction pipeline, aligned with GMAP (Wu and Nacu, 2010) to correct small InDels at splice junctions, and clustered based on intron structure or ≥95% overlap for single-exon transcripts, yielding roughly 240,000 full-length transcripts. mina- and PacBio-derived assemblies were then integrated and refined with PASA (Haas et al. 2003), producing 152,786 transcript models prior to filtering.

The genome was soft-masked using RepeatMasker (Smit et al., 2013) with a species-specific repeat library. This library combined de novo repeats predicted by RepeatModeler2 (Flynn et al., 2020) from the *R. argutus* and *R. ulmifolius* genomes, together with common Viridiplantae and Embryophyta repeats from RepBase (Bao et al., 2015) and Dfam (Hubley et al., 2016). Putative gene loci were inferred from transcript alignments and protein homology. Proteins from 24 diverse plant species (*Arabidopsis thaliana*, *Beta vulgaris*, *Cannabis sativa*, *Carya illinoinensis*, *Cucumis sativus*, *Fragaria vesca*, *Glycine max*, *Gossypium raimondii*, *Liriodendron tulipifera*, *Malus domestica*, *Medicago truncatula*, *Mimulus guttatus*, *Morus notabilis*, *Oryza sativa*, *Populus trichocarpa*, *Potentilla anserina*, *Prunus persica*, *Rosa chinensis*, *Solanum lycopersicum*, *Sorghum bicolor*, *Vitis vinifera*, and *Ziziphus jujuba*), along with Swiss-Prot eukaryotic proteins (release 2022_04), were aligned to the repeat-masked genome with EXONERATE (Slater and Birney 2005), allowing extensions up to 2 kb beyond predicted gene boundaries when non-overlapping with other loci.

Gene models were predicted using multiple approaches, including FGENESH+ and FGENESH_EST (Salamov and Solovyev, 2000), EXONERATE, AUGUSTUS (Stanke et al. 2006), and PASA-derived homology-constrained ORFs. AUGUSTUS was trained on high-confidence PASA-derived ORFs and intron hints from short read alignments. At each locus, the highest scoring prediction was selected based on transcript and protein support while avoiding predictions overlapping with repeats. PASA further refined models by adding UTRs, alternative transcripts, and correcting splice junctions.

Gene model proteins were evaluated for homology against the reference proteomes to calculate Cscores (the ratio of a protein’s BLASTP score to that of its mutual best hit) and protein coverage (percentage aligned to best homolog). Transcripts were retained if they were supported by transcripts derived from Iso-Seq and RNA-Seq or had Cscore and coverage ≥0.5. Stricter thresholds (Cscore ≥0.9 and coverage ≥70%) were applied for models with >20% CDS overlap with repetitive elements. Proteins were also annotated for domains using Pfam (Mistry et al. 2021), and models with >30% overlap with TE-related domains or lacking transcript/homology support were removed. Additional manual curation excluded incomplete or weakly supported gene models, short single-exon models (CDS <300 bp) without recognizable domains or expression evidence, and repetitive models lacking strong homology support.

### Functional annotation of *R. argutus* cv. Hillquist v.1.2

All predicted peptide sequences were functionally annotated using a computational pipeline. InterProScan 5 (Jones et al. 2014) was used to identify protein domains and sequence features, which were subsequently used to assign Gene Ontology (GO) terms. Enzyme functions were predicted with E2P2 (Chae et al. 2014; Schläpfer et al. 2017) to assign EC numbers, and PathoLogic (Karp et al. 2011) was employed to map proteins to metabolic pathways. Eukaryotic Orthologous Groups (KOG) classifications were assigned using a modified mutual best-hit algorithm. Finally, protein domain annotations were integrated to generate putative gene functional assignments, including the multiplicity of each function across the proteome.

### Whole-genome sequencing of PF and FF genotypes

A set of 17 diverse tetraploid blackberry genotypes was selected for whole-genome resequencing with the aim of identifying functional polymorphisms (SNPs and small InDels) responsible for the PF trait. Young leaf tissue of the PF genotypes Black Magic^TM^, Prime-Ark^®^ ‘45’, Prime-Ark^®^ ‘Freedom’, ‘S213’, ‘S214’ and ‘S242’, and the- FF genotypes ‘Apache’, ‘Kiowa’, ‘Osage’, ‘Ouachita’, ‘Natchez, ‘Navaho’, ‘Tupy’, ‘S9’, ‘S17’, ‘S28’ and ‘S334’ was collected on ice and stored at -80 °C until DNA isolation. Sequencing and raw data analysis were previously described by Johns et al. (2025). Briefly, 150 bp paired-end reads were cleaned and trimmed with Trimmomatic (Bolger et al. 2014) and aligned to the *R. argutus* cv. ‘Hillquist’ v.1.2 reference genome using bwa-mem2 software (Vasimuddin et al. 2019). Genetic variants were called using GATK using the option ploidy (-ploidy 4) of the “HaplotypeCaller” function for tetraploid calling. Finally, functional annotation of genetic variants was performed using NGSEP (Duitama et al. 2013), based on the GFF3 file containing gene functional annotations for the ‘Hillquist’ v.1.2 genome.

### Identification of primocane-fruiting candidate genes

To identify candidate genes responsible for the PF trait, linkage disequilibrium (LD) was estimated between SNPs on chromosome Ra03 using the ldsep R package (Gerard 2021). Based on the observed slow rate of LD decay, a genomic region surrounding the most significant PF-associated SNP was selected for further comparison between PF and FF genotypes. Genes annotated in the *R. argutus* cv. ‘Hillquist’ v1.2 genome within this region with a gene function associated with flowering were considered candidate genes. Candidate genes were further evaluated using whole-genome sequencing data to identify SNPs and InDels predicted to alter the encoded protein between PF and FF genotypes.

### KASP marker development and validation

Whole-genome sequence data were used to identify polymorphisms within the genomic region associated with the PF trait in the GWAS and genetic linkage analysis. These polymorphisms were then used to develop diagnostic Kompetitive Allele-Specific PCR (KASP) markers for predicting the PF trait. Because the WGS data were aligned to the ‘Hillquist’ reference genome, the original donor of the PF trait, and it has been reported that a single recessive allele controls the PF trait (Lopez-Medina et al. 2000), SNPs were selected based on genotype patterns consistent with this inheritance. In the six PF genotypes, selected SNPs had to be homozygous for the reference allele (AAAA, where “A” denotes the reference allele), whereas in the 11 FF genotypes, the selected SNPs were required to be heterozygous or homozygous for the alternative allele (AAAB, AABB, ABBB, or BBBB, where “B” represents the alternative allele). Twelve SNPs located within 1 Mb of the most significant GWAS peak were selected as targets for KASP assay design by LGC Genomics (Beverly, MA, USA). Each KASP assay contains two allele-specific forward primers, each with a unique tail sequence serving as a binding site for oligonucleotides labeled with fluorescent dyes FAM or HEX, and one common reverse primer.

To evaluate the accuracy of the KASP markers to predict the PF trait, a set of 494 genotypes with diverse genetic backgrounds from the UADA program (including 165 seedlings from the biparental population 1937 and 201 cultivars and selections), the United States Department of Agriculture – Horticultural Crops Production and Genetic Improvement Research Unit (USDA-ARS-HCPGIRU, n=76), Hortifrut Genetics (HFG, n=39), and the United States Department of Agriculture – National Clonal Germplasm Repository (USDA-ARS-NCGR, n=13), was genotyped at LGC Genomics (Beverly, MA, USA). A total of 167 out of the 367 genotypes included in the GWAS panel were also represented in the validation panel, comprising 155 cultivars and selections from UADA, two cultivars from USDA-ARS HCPGIR, and ten accessions from USDA-ARS-NCGR. KASP genotyping was performed by LGC Biosearch Technologies™ as described in Johns et al. (2025). To determine allele dosage, the fluorescence values measured for the FAM and HEX dyes for each KASP marker were analyzed using SNPviewer software (Biosearch Technologies™). Final cluster plots were generated using the ggplot R package (Wickham, 2016).

## Results

### GWAS Reveals a Major Locus on Chromosome Ra03 Controlling the PF Trait

A set of 137 PF and 228 FF blackberry genotypes were used in the genome-wide association study (Supplementary Table 1). Of the 124,564 SNPs initially identified in a larger panel of 495 blackberry genotypes, 81,064 biallelic SNPs distributed across the seven blackberry chromosomes were retained for association analysis in 365 genotypes. The final set of markers was obtained after filtering those with a maximum genotype frequency higher than 1 – 5/N, where N represents the population size, as recommended for heterozygous panels by Rosyara et al. (2016). Chromosome-specific QQ-plots revealed an upward curve only on chromosome Ra03, indicating no technical biases or population structure affecting the association (Supplementary Fig. 1). Using the ‘M.eff’ method to control the genome-wide false positive rate, 419 significant SNPs associated with the PF trait were detected within a genomic region located on chromosome Ra03 between 21,158,993 and 41,668,169 bp (Supplementary Table 2). The most strongly associated SNPs were located at 33,338,602 bp and 33,338,650 bp, each with a - log_10_(p) of 183.8 under the simplex dominant (1-dom-ref) model (Fig. 2). Both variants are located within an intron of the gene Ruarg.3G335600, which is homologous to the gene AT2G28470 that encodes the enzyme Beta-galactosidase 8 (BGAL8) in *A. thaliana*. For both SNPs, 133 out of 137 PF blackberry genotypes (97%) were homozygous for the reference allele (the allele associated with the presence of the PF trait). In contrast, all FF genotypes were either homozygous for the alternative allele or heterozygous.

**Figure 2.**
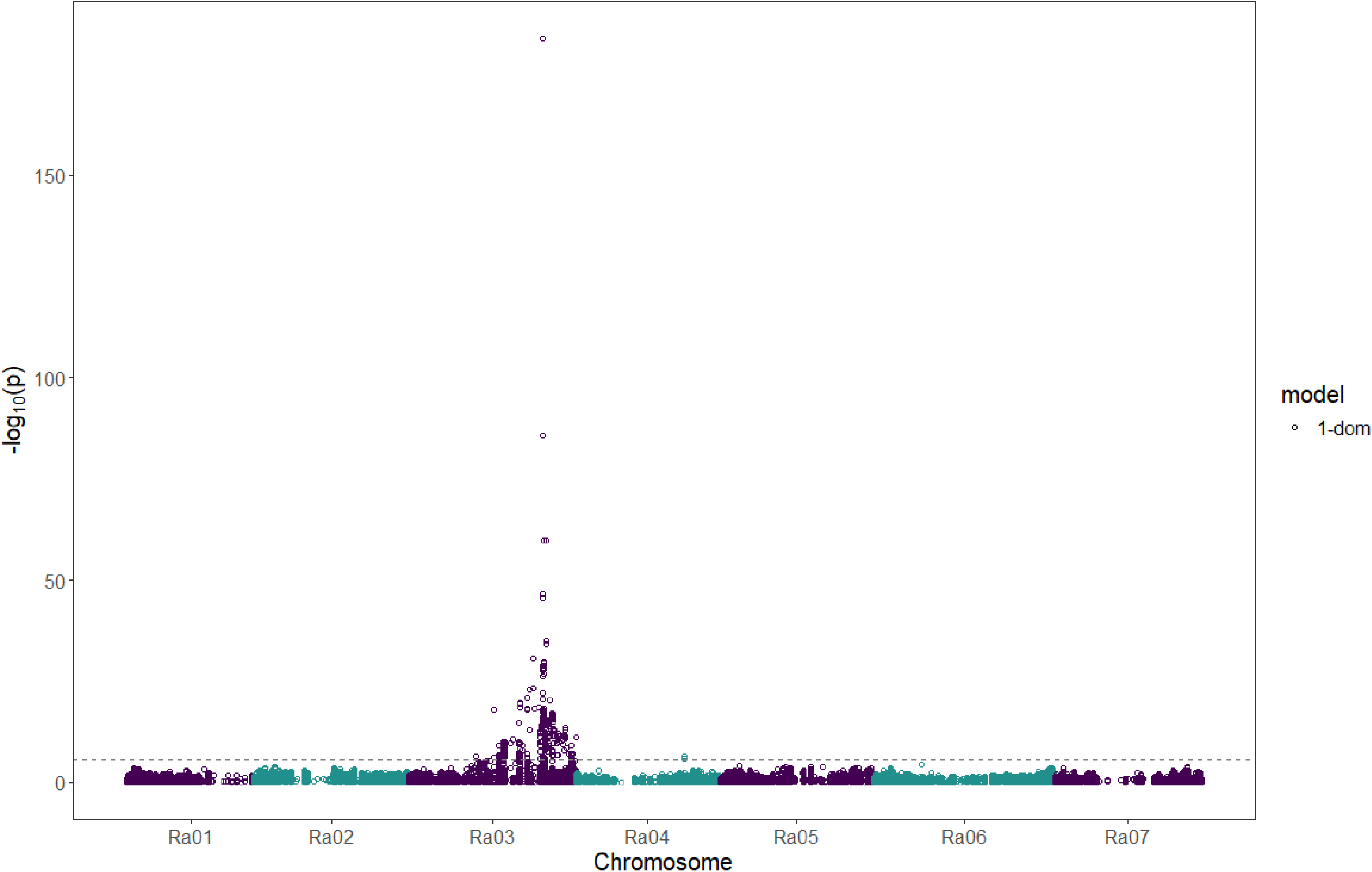
Manhattan plot for the association analysis between Primocane-Fruiting trait and SNPs under the simplex-dominant (1-dom) model. The dashed line represents the-log10(p) threshold using the “M.eff” method implemented in GWASpoly.

### Genetic Linkage Mapping Supports Placement of PF locus on Chromosome Ra03

The first integrated SNP-based linkage map in blackberry was constructed to identify loci associated with the PF trait. A total of 165 seedlings derived from the cross between the FF selection ‘S16’ and the PF selection ‘S242’ were evaluated for the presence of flowering primocanes during two consecutive years and submitted to Allegro^®^ targeted genotyping. Ninety progenies were recorded as PF, 72 as FF, and three were undetermined. Thirty-six progenies with a low number of Allegro® targeted genotyping reads were excluded from the linkage analysis. After filtering out SNPs with low quality, 9,736 SNP markers were initially selected to construct the genetic linkage map (Supplementary Table 3). The most strongly associated SNP markers identified in the GWAS (Ra03:33,338,650 and Ra03:33,338,602) were also evaluated in the 1937 biparental population (see Development and Validation of KASP Markers section).

However, Ra03:33,338,650 was excluded from linkage mapping due to missing data in 13 out of the 113 progenies (more than 10%). A Principal Component Analysis (PCA) was performed to identify potential self-cross individuals or contaminants in the biparental population. As a result, 16 individuals were identified as potential non-true F_1_ samples and were excluded from the analysis (Supplementary Fig. 2). Therefore, the linkage map was constructed using 113 F_1_ seedlings, of which 62 were recorded as PF, 50 were recorded as FF, and one was undetermined.

Because both ‘S16’ and ‘S242’ were included in the GWAS panel, allele dosages for the most significant markers linked to the PF trait could be estimated for both parents. The FF female parent, ‘S16’, carries three copies of the reference allele associated with the PF trait, as indicated by genotypes at markers Ra03:33,338,650_G/A (GGGA) and Ra03:33,338,602_T/C (TTTC). The PF male parent, ‘S242’, was homozygous for the PF allele at both loci, with genotypes GGGG and TTTT, respectively (Supplementary Table 1). Assuming segregation from a triplex (three PF alleles) × quadriplex (four PF alleles) cross, the observed PF:FF segregation ratio of the offspring fit the expectation for a tetrasomic inheritance model for a trait controlled by a single recessive allele, under both random chromosome assortment (expected ratio 1:1; χ² = 1.286; P = 0.257) and random chromatid assortment (expected ratio 15:13; χ² = 0.144; P = 0.705).

After filtering markers with >10% missing data, monomorphic loci, redundant markers, and those showing significant segregation distortion (χ² test, P < 0.05), a final set of 3,882 SNPs was retained for linkage mapping using MAPpoly2. Pairwise recombination fractions were computed, and the resulting matrix exhibited the expected seven block-diagonal clusters corresponding to the seven blackberry linkage groups (LGs) (Fig. 3a). The genetic map was assembled assuming a 5% global error rate and using the ‘Hillquist’ reference genome to guide marker order (Mollinari and Garcia, 2019). The final map spanned 769.4 cM and included 3,197 non-redundant SNPs, averaging four markers per cM across LGs ranging from 86.5 cM (LG4) to 122.2 cM (LG2) (Fig. 3b, Table 1). Marker density was highest in LG2 (618 SNPs; 5.06 markers/cM) and lowest in LG4 (208 SNPs; 2.41 markers/cM), with the largest gaps observed in LG4 (14.7 cM) and LG5 (5.3 cM). The map was highly collinear with the ‘Hillquist’ physical map, showing no apparent inversions or translocations (Supplementary Fig. 3). Probability profiles of all meiotic pairing configurations indicated no evidence of preferential pairing in either parent (Supplementary Fig. 4).

**Figure 3.**
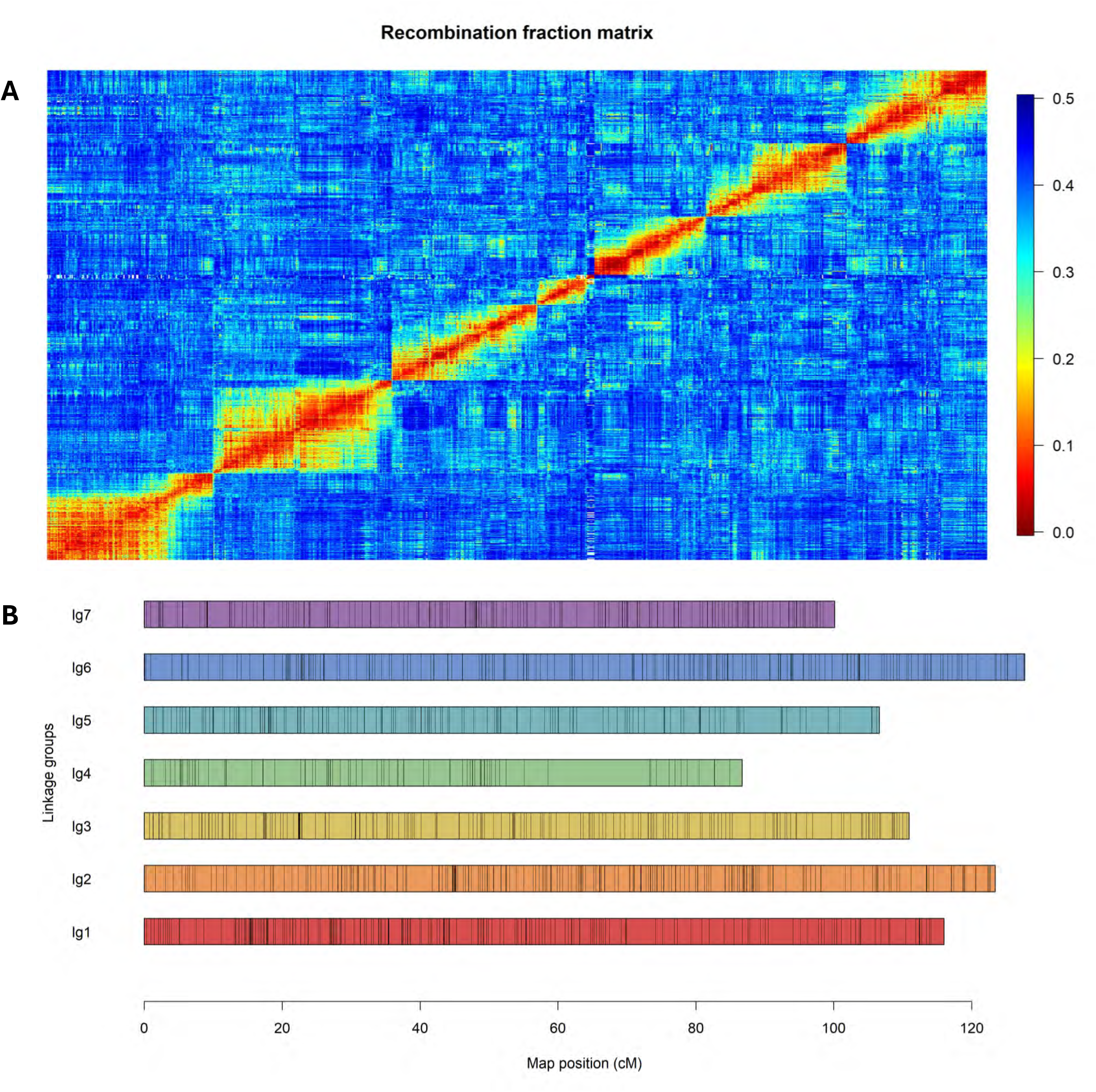
Linkage map construction for the blackberry biparental population 1937 derived from the cross between the FF genotype ‘S16’ and the PF genotype ‘S242’. (A) Recombination fraction matrix showing seven submatrices corresponding to each of the seven linkage groups (basic chromosome number in blackberry). (B) Integrated linkage groups (LGs) constructed by combining the individual genetic maps from both parents.

**Table 1.** Genetic map summary of the tetraploid blackberry biparental population 1937, derived from a cross between the Floricane-fruiting (FF) genotype ‘S16’ and Primocane-fruiting (PF) genotype ‘S242.’

| LG | Map length<br>(cM) | Markers/cM | Simplex_P1 | Simplex_P2 | Double-<br>simplex | Multiplex | Total | Max gap |
| --- | --- | --- | --- | --- | --- | --- | --- | --- |
| 1 | 115.9 | 4.94 | 234 | 40 | 95 | 204 | 573 | 4.3 |
| 2 | 122.2 | 5.06 | 26 | 26 | 268 | 298 | 618 | 3.7 |
| 3 | 110.5 | 4.24 | 53 | 86 | 103 | 226 | 468 | 3.3 |
| 4 | 86.5 | 2.41 | 44 | 52 | 39 | 73 | 208 | 14.7 |
| 5 | 106 | 3.52 | 61 | 10 | 46 | 256 | 373 | 5.3 |
| 6 | 128 | 3.91 | 66 | 41 | 165 | 228 | 500 | 3.1 |
| 7 | 110.3 | 4.56 | 46 | 72 | 144 | 195 | 457 | 3.2 |
| Total | 769.4 | 4 | 530 | 327 | 860 | 1480 | 3197 | 14.7 |

The presence or absence of the PF phenotype in the biparental population was included as a phenotypic marker in the linkage map and positioned at 75.3 cM on linkage group 3 (LG3) (Fig. 4). The PF-linked marker Ra03:33,338,602 correctly classified all progeny exhibiting the PF trait as quadruplex (‘TTTT’), except for four individuals (1937-79, 1937-90, 1937-129, and 1937-141), which were predicted to be triplex (‘TTTC’). This marker was also positioned at 75.3 cM on LG3. The recombination fraction between each molecular marker and the PF phenotype analysis revealed a maximum LOD score of 33.7 for the marker Ra03:33,338,602. This was followed by the markers Ra03:32,702,539, Ra03:31,379,074 and Ra03:34,733,921, which had LOD scores of 23.1, 22.1 and 20.9, respectively (see Fig. 4a and Supplementary Table 4). These results, combined with the GWAS analysis, suggest that the PF trait is controlled by a major recessive locus between 31 and 35 Mb on chromosome Ra03.

**Figure 4.**
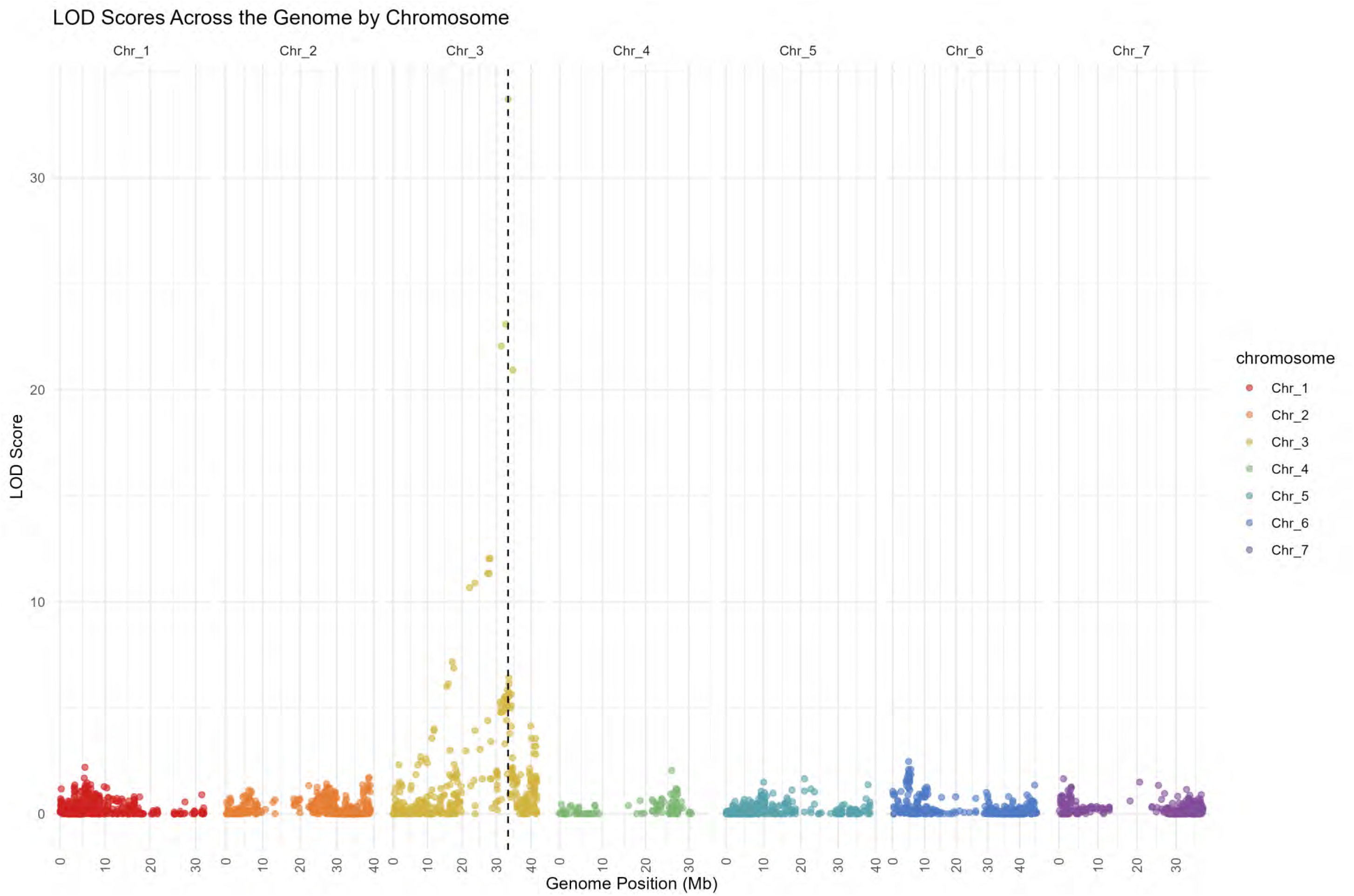
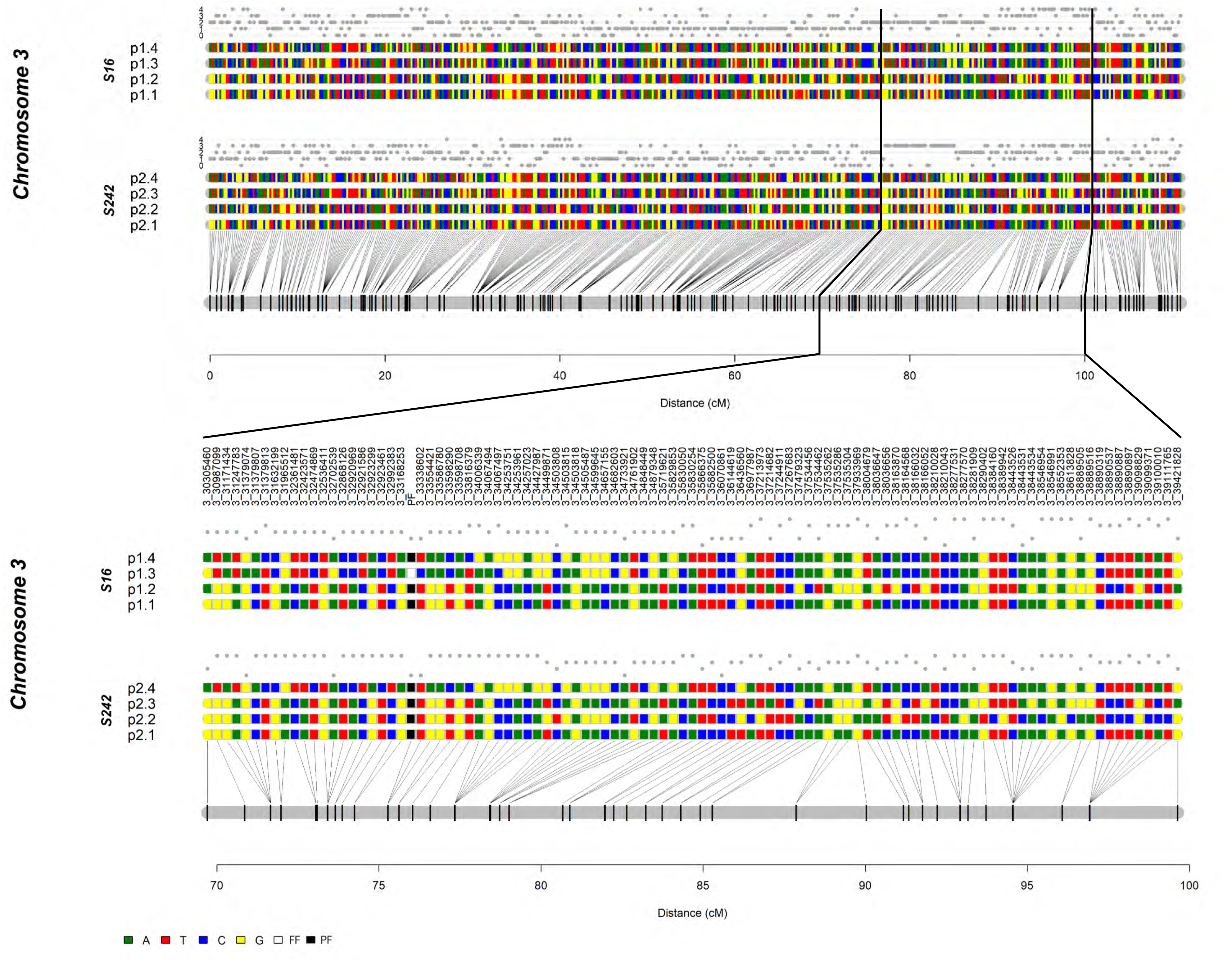
(A) Recombination fraction calculation between the PF phenotype in the biparental population (presence or absence of the PF trait) and each molecular marker in the biparental population ‘1937’. The dashed line indicates the position of the maximum LOD score found on the chromosome 3 (Ra03). (B) Linkage genetic analysis of LG3 (upper panel) with a zoom in the region between 70 and 100 cM showing the haplotype for the SNPs linked to the marker PF1 (Ra03:33,338,602) and to the PF locus. The colored rectangles indicate SNP nucleotides within each of the four haplotypes for S16 (p1.1 to p1.4) and for S242 (p2.1 to p2.4). White and black rectangles indicate the FF and PF phenotypic allele. The allele dosage is shown above each parent’s homologs.

### Updated annotation of *R. argutus* cv. ‘Hillquist’

Before structural annotation, 126 Mb (42.3%) of the ‘Hillquist’ genome was repeat-masked. The final set of predicted genes contained 30,026 coding genes, corresponding to 43,850 coding transcripts when alternative isoforms were included (Table 2). The 30,026 coding genes (calculated from the primary isoform of each gene) had an average length of 1,239 bp and a mean exon number of 5.04. Of these coding genes, 21,843 had no alternative isoforms, 5,068 had two isoforms, and 3,115 had three or more isoforms. In the predicted set of genes, 2,149 (92.4%) complete *R. argutus* genes orthologous to the eudicots_odb10 BUSCO families were identified, along with 76 (3.3%) genes with partial matches. A small fraction of the BUSCO families (4.3%) was not identified among the predicted *R. argutus* genes in the updated annotation (Table 2).

**Table 2.** Comparison of the original and updated *R. argutus*’Hillquist’ genome annotations. *Calculated from the primary isoform of each gene.

|  |  | v.1 (Bruna et al. 2023) | v.1.2 |
| --- | --- | --- | --- |
| Gene prediction summary | Primary transcripts (loci) | 38,503 | 30,026 |
|  | Genes with more than 1 transcript isoform | 1,667 | 8,183 |
|  | Mean CDS length (bp)* | 1,021 | 1,239 |
|  | Median coding exon length (bp)* | 132 | 135 |
|  | Mean count of coding exons per gene* | 4.36 | 5.04 |
| BUSCO completedness (eudicots_odb10, n= 2,326) | Complete (C) | 91.70% | 92.40% |
|  | Fragmented (F) | 3.20% | 3.30% |
|  | Missing (M) | 5.10% | 4.30% |
| Functional annotation | Genes with PFAM domain (% of all genes) / unique PFAM domains | 24,859 (65.6%) / 4,324 | 23,976 (79.9%) / 4,359 |
|  | Genes with PANTHER domain (% of all genes) / unique PANTHER domains | 28,497 (74.0%) / 6,243 | 27,517 (91.6%) / 6,318 |
|  | Genes with KEGG pathway (% of all genes) / unique KEGG pathways | 7,468 (19.4%) / 3,290 | 7,699 (25.6%) / 3,330 |
|  | Genes with KOG classification (% of all genes) / unique KOG domains | 9,188 (23.9%) / 2,467 | 10,186 (33.9%) / 2,563 |
|  | Genes with InterPro assignment (% of all genes) / unique InterPro assignments | 27,184 (70.6%) / 6,353 | 25,517 (85.0%) / 6,420 |
|  | Genes with any functional annotation (% of all genes) | 31,847 (82.7%) | 28,881 (96.2%) |
| Support for predicted gene models | PSAURON score* | 87.9 | 95.4 |
|  | Genes supported by transcript alignments* | 23,417 | 23,840 |
|  | % of genes supported by transcript alignments* | 60.80% | 79.40% |

These results suggest that the *R. argutus* cv.‘Hillquist’ v.1.2 assembly and annotation are 95.7% complete. Evidence supporting the predicted gene models was strong. For 79.4% of loci, the coding regions of the primary transcripts were supported by Iso-Seq or RNA-seq alignments covering at least 80% of the coding length and showing ≥80% sequence identity. Additionally, 27,328 transcripts displayed >50% peptide homology to known proteins in other plant species.

Functional annotation was similarly comprehensive: 28,881 (96.2%) of predicted genes received at least one functional annotation. Pfam domains were detected in 23,976 genes (79.9%), spanning 4,359 unique Pfam families, while PANTHER domains were identified in 27,517 genes (91.6%) across 6,318 unique families. Kyoto Encyclopedia of Genes and Genomes (KEGG) pathway mappings were assigned to 7,699 genes (25.6%), covering 3,330 unique pathways, and Eukaryotic Orthologous Group (KOG) classifications were assigned to 10,186 genes (33.9%), encompassing 2,563 unique KOG domains. Collectively, these results indicate that the majority of predicted genes in the ‘Hillquist’ v1.2 annotation have putative biological functions supported by conserved domain, pathway, or orthology evidence.

### Putative candidate genes associated with the PF trait

Both GWAS and genetic linkage mapping identified a region between 31 and 35 Mb on chromosome Ra03 with peak significance at 33,338,602 bp as associated with the PF trait. Within this interval, ten genes with predicted functions related to flowering regulation were identified in the ‘Hillquist’ genome (Table 3). These include UPSTREAM OF FLC (Ruarg.3G323500), BEL1-like homeodomain 8 (BLH8; Ruarg.3G329500), Homeobox-leucine zipper protein REVOLUTA (REV; Ruarg.3G332300), and a CCCH-type zinc finger protein (Ruarg.3G334700). Additional genes in the interval include a DNA-binding protein with MIZ/SP-RING zinc finger (SIZ1; Ruarg.3G337900), AGAMOUS-like MADS-box protein AGL28 (Ruarg.3G338800), an AP2/ERF family transcription factor (Ruarg.3G343800), a ubiquitin-specific protease (UBP; Ruarg.3G346400), an AP2/B3-like transcription factor (Ruarg.3G349500), and a B3 domain-containing transcription factor VRN1 (Ruarg.3G349600).

**Table 3.** Candidate genes for the Primocane-fruiting (PF) trait in blackberry located on chromosome Ra03 between 31 and 35 Mb.

| Gene ID | Start position | End position | Distance from GWAS peak (Kb) | Predicted gene function | Arabidopsis homolog |
| --- | --- | --- | --- | --- | --- |
| Ruarg.3G323500 | 32004799 | 32009449 | 1329 | UPSTREAM OF FLC protein | AT2G28150 |
| Ruarg.3G329500 | 32620726 | 32627768 | 710 | BEL1-like homeodomain 8 | AT2G27990 |
| Ruarg.3G332300 | 32985451 | 32993007 | 346 | Homeobox-leucine zipper protein REVOLUTA | AT5G60690 |
| Ruarg.3G334700 | 33264790 | 33269811 | 68 | Zinc finger (CCCH-type) family protein | AT2G28450 |
| Ruarg.3G337900 | 33596729 | 33608051 | 258 | DNA-binding protein with MIZ/SP-RING zinc finger, PHD-finger and SAP domain-containing protein | AT5G60410 |
| Ruarg.3G338800 | 33668587 | 33669717 | 330 | Agamous-like MADS-box protein AGL28 | AT1G01530 |
| Ruarg.3G343800 | 34253342 | 34257478 | 915 | Ethylene-responsive transcription factor RAP2-7 (Related to AP2.7) | AT2G28550 |
| Ruarg.3G346400 | 34566835 | 34572407 | 1228 | Ubiquitin specific protease domain protein | AT5G06600 |
| Ruarg.3G349500 | 34872157 | 34874071 | 1533 | AP2/B3-like transcriptional factor family protein | AT1G49475 |
| Ruarg.3G349600 | 34879197 | 34881066 | 1540 | B3 domain-containing transcription factor VRN1 | AT3G18990 |

### Genetic variants within the genomic region associated with the PF trait

The whole-genome resequencing data from 17 diverse tetraploid blackberry genotypes were analyzed to identify SNPs and small InDels in the genomic region on chromosome Ra03 associated with the PF trait, with the aim of pinpointing the causal variant. Genetic variants within and adjacent to potential candidate genes associated with the PF trait were first investigated. Additional variants located within 1 Mb downstream and upstream from the most significant SNP associated with the PF trait (Ra03:33,338,602) were also investigated. A total of 151,854 variants were initially detected after quality filtering in this genomic window. Because previous genetic studies (Castro et al. 2013; Lopez-Medina et al. 2000) and our GWAS results support a single recessive allele model for the PF trait, we prioritized variants where PF genotypes were homozygous for the reference allele, while FF genotypes were heterozygous or homozygous for the alternative allele. A total of 607 SNPs and small InDels were identified that fit this criterion. Of these variants, 24 were located within or adjacent to four of the ten candidate genes in the region with putative functions related to flowering (Table 4). One small InDel located at 32,625,674 bp was identified in an intron of the gene Ruarg.3G329500, which encodes for a BEL1-like homeodomain 8 (BLH8). Two SNPs and one InDel variant were identified in the intron region and one InDel within 1 Kb upstream of the gene Ruarg.3G332300, which codes for a Homeobox-leucine zipper protein REVOLUTA. Twelve variants were identified within and adjacent to the CCCH-type Zinc finger family protein-coding gene (Ruarg.3G334700), including three intron variants, one synonymous SNP in an exon region, two 3’UTR variants, and six variants within 1 Kb downstream of the gene. Furthermore, seven variants were found within or adjacent to the ubiquitin-specific protease (Ruarg.3G346400), including two intron variants, three SNPs in the 3’UTR region and two SNP within 1 Kb downstream of the gene.

**Table 4.**
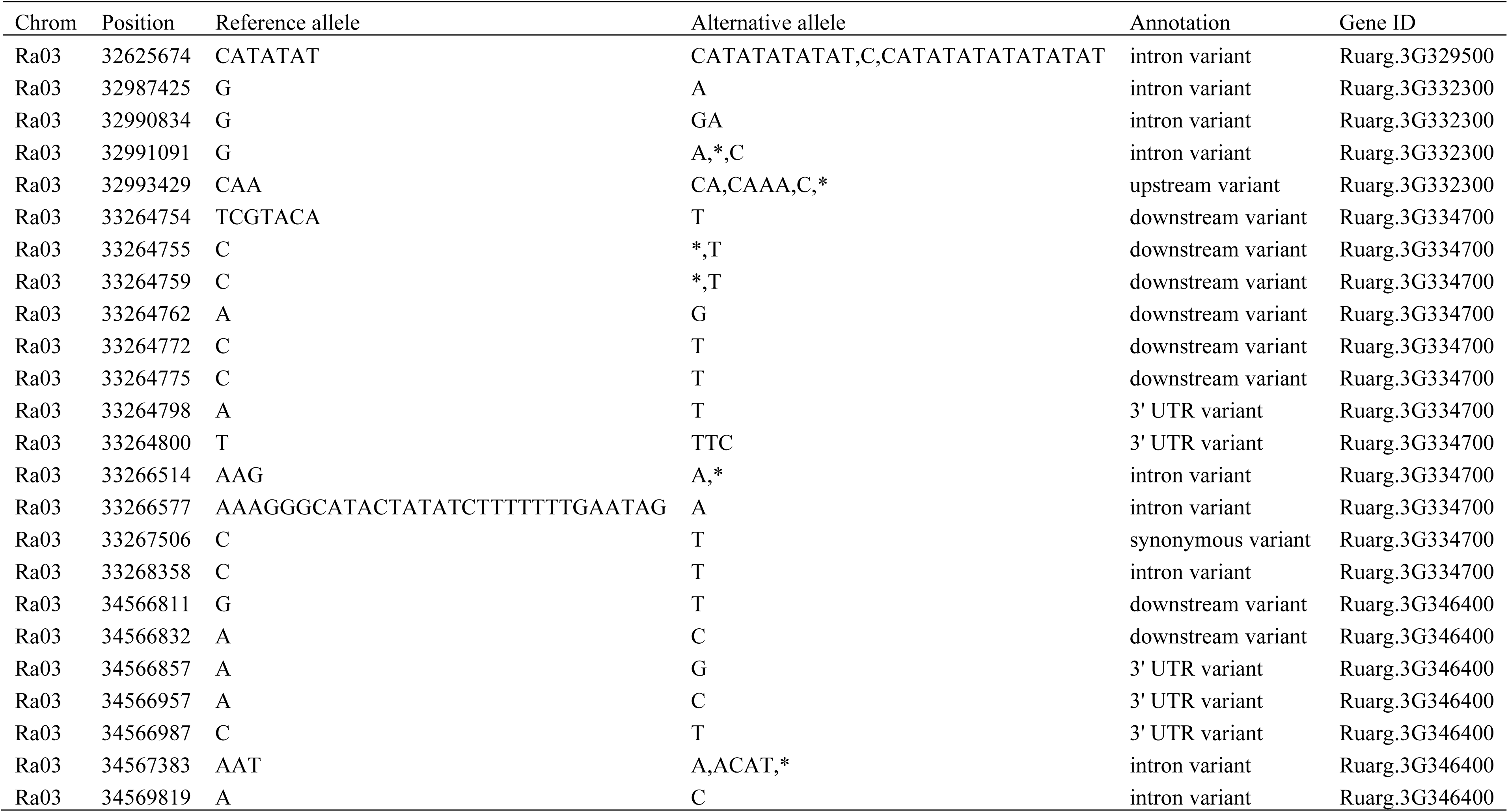
Variants identified within and adjacent to potential candidate genes in the primocane-fruiting (PF) locus in blackberry, with functional annotations related to flowering development, based on whole-genome sequencing data from 17 blackberry selections and cultivars. These variants distinguish PF (six genotypes) from FF (11 genotypes) based on the WGS data. Asterisk represents a deletion spanning over the variant position.

The additional 583 genetic variants detected within the 2 Mb window around the GWAS peak were all located in either intergenic regions or within non-candidate genes (Supplementary Table 5). From this set of variants, 12 SNP markers were selected for KASP development; the two most strongly associated with the PF trait in the GWAS located at 33,338,602 and 33,338,650 bp, and ten other SNPs that discriminate between the PF and FF genotypes at positions 33,713,953 bp, 33,968,573 bp, 34,098,961 bp, 34,100,366 bp, 34,100,510 bp, 34,106,515 bp, 34,287,111 bp, 34,352,079 bp, 34,364,423 bp and 34,399,952 bp (Supplementary Table 6 and 7).

### Development and validation of diagnostic KASP markers

Among the 12 KASP markers, two were identified as the most predictive in the validation panel. These two markers, targeting the SNPs located at 33,338,650 and 33,338,602 bp on chromosome Ra03, named *PF1* and *PF2*, respectively, predicted the PF phenotype with 96.8% and 96.5% accuracy, in the validation panel (Fig. 5 and Supplementary Table 8). These SNPs were the markers most strongly associated with the PF trait in the GWAS analysis (Fig. 2 and Supplementary Table 2).

**Figure 5.**
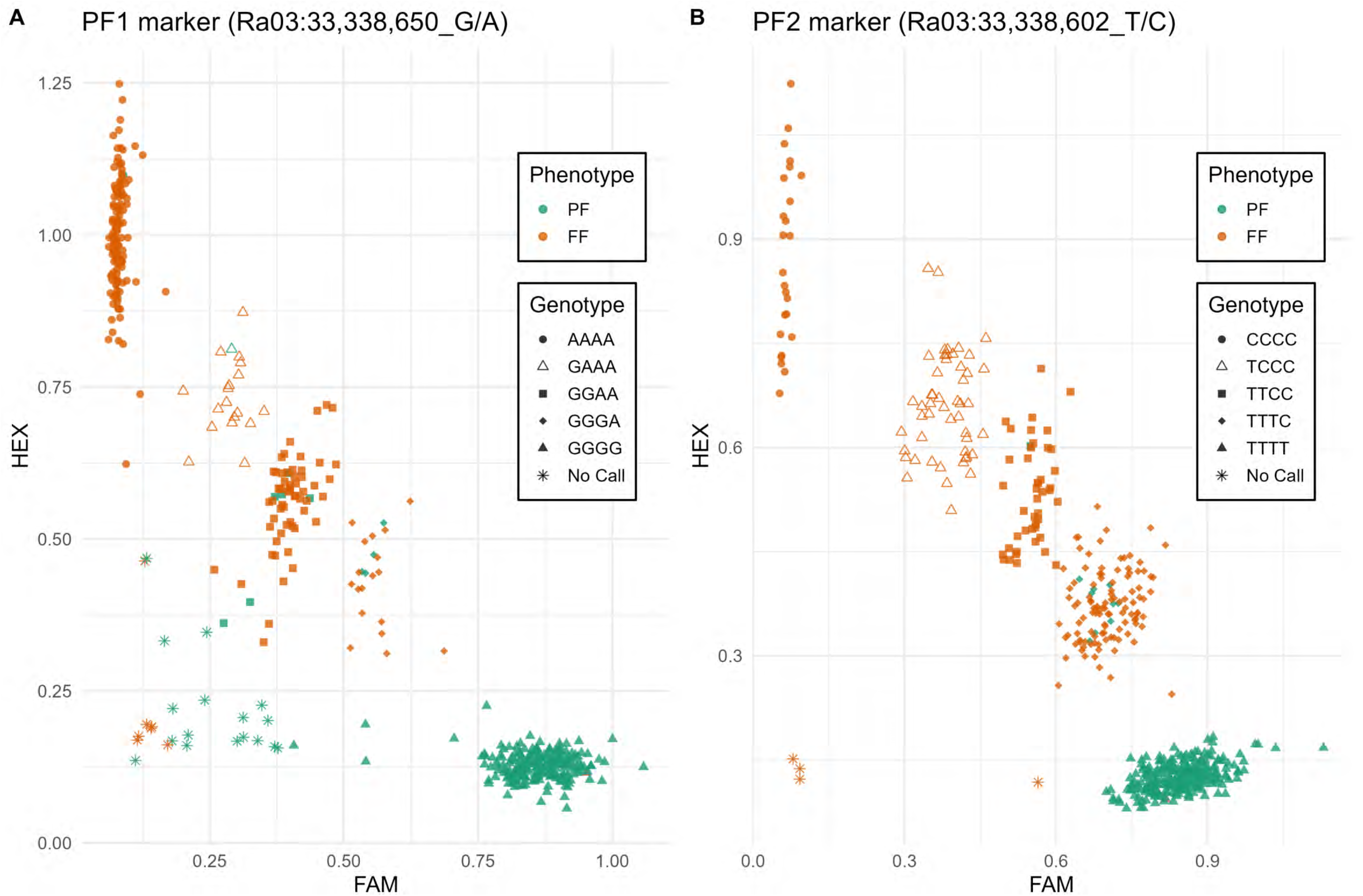
Validation of KASP markers. The scatter plot displays the FAM and HEX values for 647 genotypes for marker PF1 (A), which targets the SNP at Ra03: 33,338,650 (G/A), and marker PF2 (B), which targets the SNP at Ra03: 33,338,602 (T/C). The allele dosage for the simplex, duplex, and triplex classes was determined based on the fluorescence intensity of the FAM and HEX dyes.

The *PF1* and *PF2* marker reactions failed to call alleles for 24 and 4 of 494 genotypes in the validation panel, respectively. Among the genotypes with successful allele calls, 455 and 473 were correctly predicted by *PF1* and *PF2* markers, respectively. Both markers incorrectly predicted 11 genotypes. Five seedlings from population 1937 (1937-001, 1937-079, 1937-090, 1937-129, and 1937-141), three selections from the UADA breeding program (S307, S441, and S443), and two selections from Hortifrut Genetics Ltd. (HFG39 and HFG45) were all scored as PF genotypes but had at least one non-PF allele at both *PF1* and *PF2* sites (Supplementary Table 8). On the other hand, one seedling from population 1937 (1937-029) was scored as a FF genotype but was predicted to be homozygous for the PF allele at both *PF1* and *PF2* markers.

Six additional genotypes were inaccurately predicted by only one of the two markers. One selection from the USDA-ARS HCPGIR germplasm (OR72) and one from the UADA breeding program (S306) were scored as PF genotypes but were predicted to have only two PF alleles for the *PF1* marker. Finally, one seedling from population 1937 (1937-107), one selection from the UADA program (S140), and two selections from the USDA-ARS HCPGIR germplasm (OR34 and OR53) were scored as FF genotypes but were predicted to have only PF alleles for the *PF2* marker.

## Discussion

### A single major locus on chromosome Ra03 controls PF in tetraploid blackberries

Genome-wide association analysis across 365 tetraploid blackberry genotypes revealed a single major locus on chromosome Ra03 strongly associated with the PF trait. Using 81,064 high-quality biallelic SNPs distributed across the seven blackberry chromosomes, 419 SNPs were significantly associated with PF within a 20.5 Mb region on Ra03 (21.2–41.7 Mb), with the most strongly associated variants located at 33,338,602 and 33,338,650 bp (−log₁₀P = 183.8). Nearly all PF genotypes (97%) were homozygous for the reference allele at this locus, while all FF genotypes had at least one copy of the alternative allele. This clear segregation pattern indicates that a single, major-effect recessive locus underlies PF in tetraploid blackberry.

The GWAS findings were independently validated through linkage analysis in a biparental F₁ population derived from a cross between the FF selection ‘S16’ and the PF selection ‘S242’. The PF phenotype mapped to linkage group 3 (LG3) at 75.3 cM and was tightly linked to a KASP marker (*PF2*) targeting the peak GWAS SNP Ra03:33,338,602. The maximum LOD score for association with PF was observed at marker Ra03:33,338,602 (LOD = 33.7), followed by Ra03:32,702,539 (LOD = 23.1), Ra03:31,379,074 (LOD = 22.1) and Ra03:34,733,921 (LOD = 20.9). Taken together, the GWAS and linkage results indicate that PF in blackberry is controlled by a major recessive locus located between 31 and 35 Mb on chromosome Ra03.

Earlier work by Castro et al. (2013) placed the PF locus between SSR markers RH_MEa0006aC04-175 and RH_MEa0007aG06-152, which were initially assigned to linkage group 7 but later shown to align to chromosome Ra02 of the ‘Hillquist’ genome assembly. No associations with the PF trait were detected on Ra02 or Ra07 in the present study, suggesting that the earlier placement may have resulted from limited marker density and the absence of a reference genome. In contrast, the current study used 3,197 SNP markers to construct a high-density linkage map spanning 769.4 cM across seven chromosomes, providing substantially higher resolution and collinearity with the reference genome. Unlike previous studies limited to single crosses, our approach also captures the genetic diversity and allelic structure present in breeding germplasm, enabling more accurate localization of the PF locus and identification of tightly linked diagnostic SNPs suitable for marker-assisted selection.

The PF mapping results in blackberry contrast with those reported for red raspberry, where Jibran et al. (2019) identified two quantitative trait loci, RiAF3 and RiAF4, located on chromosomes 3 and 4 of *R. idaeus*. These regions correspond to chromosomes Ra03 (0–9 Mb) and Ra04 (31–32 Mb) of the *R. argutus* cv. Hillquist genome, neither of which overlapped the PF locus identified here. The lack of shared loci or candidate genes suggests that primocane fruiting in tetraploid blackberry and red raspberry may have evolved through distinct genetic mechanisms. Together, the GWAS and linkage mapping results firmly establish the PF locus on chromosome Ra03 as the major determinant of annual flowering in tetraploid blackberry.

### Additional insights from high-density linkage mapping in blackberry

We constructed the first integrated high-density genetic linkage map for tetraploid blackberry using 113 progeny from a cross between the FF selection ‘S16’ and the PF selection ‘S242’. The final map spanned 769.4 cM and included 3,197 nonredundant SNP markers, averaging approximately four markers per cM across seven LGs corresponding to the basic chromosome number in blackberry. All seven linkage groups exhibited high collinearity with the *R. argutus* cv. ‘Hillquist’ reference genome (Supplementary Fig. 3), confirming the accuracy of marker ordering and alignment. The fewest markers and largest gap were observed on LG4 (Ra04), consistent with reduced recombination and genetic diversity in the distal region of this chromosome. This region coincides with the prickle-free locus, which has been shown to contain an extensive linkage disequilibrium block (Chizk et al. 2023; Johns et al. 2025). Because both parents of the mapping population are prickle-free, the paucity of markers on Ra04 may reflect historical selection and a resulting loss of heterozygosity in this region.

This map represents a substantial advance in marker density and genomic coverage compared with the first SSR-based blackberry map of Castro et al. (2013), which comprised only 119 markers and lacked the resolution to anchor linkage groups to specific chromosomes. It also improves upon the pseudo-testcross maternal map of the breeding selection ‘A-2551TN’ (Brůna et al. 2023) by incorporating markers across all dosage classes and producing a fully integrated representation of all four homologs for each chromosome, rather than relying solely on simplex markers to generate haplotype-resolved but non-integrated linkage groups.

No evidence of preferential pairing among homologs was detected in either parent (Supplementary Fig. 4), suggesting predominantly polysomic inheritance in tetraploid blackberry. In polyploid species, double reduction can occur when multivalent pairing causes sister chromatid segments to migrate into the same gamete during meiosis (Mather 1936; Bourke et al. 2015), thereby increasing homozygosity and altering expected segregation ratios. In this mapping population, the FF female parent ‘S16’ had three copies of the PF allele (TTTC) at SNP marker Ra03:33338602 (*PF2*), while the PF parent ‘S242’ had four copies of the PF allele (TTTT). A single FF progeny (1937-131) exhibited the duplex genotype (TTCC) for the *PF2* locus. Previously, Vaughn et al. (2023) reported six progenies with the duplex genotype in this population; however, five were excluded from the linkage map as non-true seedlings in the PCA analysis, most likely resulting from pollen contamination or self-pollination (Supplementary Fig. 2). The remaining progeny, 1937-131, is most likely an actual F_1_ seedling of the ‘S16’ x ‘S242’ cross, providing possible evidence of low-frequency double reduction in this mapping population. Collectively, these results establish a robust, genome-aligned linkage framework for tetraploid blackberry that captures the full complexity of polysomic inheritance and provides a foundation for future trait mapping and comparative genomic analyses.

### An improved annotation of *R. argutus* cv. ‘Hillquist’

The updated *R. argutus* ‘Hillquist’ annotation (v1.2) represents a substantial improvement over the original annotation, both in gene model accuracy and functional characterization. The final set of 30,026 coding genes is significantly smaller than the original 38,503, reflecting more stringent filtering of likely false predictions: the original annotation included 9,407 genes with no transcriptomic or protein homology support. Despite fewer predicted genes, the updated annotation exhibits higher completeness and confidence, with more BUSCO eudicot markers recovered, better transcriptome support (both in relative and absolute terms) and more comprehensive functional annotation. In particular, compared to the original annotation, the updated version shows consistently higher proportions of genes assigned to Pfam, PANTHER, KEGG pathways, and KOG categories, as well as a large increase in the percentage of genes receiving any functional annotation (82.7% versus 96.2%). These improvements are accompanied by increases in the number of unique domains or pathways captured across Pfam, PANTHER, KEGG, and KOG classifications, reflecting broader and more diverse functional coverage (Table 2). Together, these gains indicate that the improved structural annotation is matched by richer and more reliable biological characterization. Finally, the updated annotation achieved a markedly better Protein Sequence Assessment Using a Reference ORF Network (PSAURON) score, with PSAURON providing a genome-wide measure of protein-coding sequence accuracy using a machine learning model trained across diverse taxa (Sommer et al. 2025).

Several methodological improvements contributed to these gains. Repeat masking was refined by manually curating predicted transposable element libraries to prevent genuine genes from being erroneously masked. The annotation pipeline also fully integrated long-read Iso-Seq data, rather than using it solely for validation of gene models and minor repeat-masking refinement in the original annotation. This integration enabled correction of mispredicted gene models and improved gene prediction in regions lacking homology to known proteins. Furthermore, the set of protein homologs used for functional annotation was curated to include additional close relatives, improving the accuracy of homology-derived models. The updated annotation also includes UTR predictions, which may be particularly relevant for functional analyses of regulatory variants, and a significantly larger set of predicted alternative isoforms. Overall, these improvements provide a more accurate and biologically meaningful representation of the *R. argutus* gene space, supporting downstream analyses including candidate gene identification and variant interpretation for traits such as PF.

### Putative Candidate Genes for PF

The PF locus on chromosome Ra03 was defined by both GWAS and genetic linkage mapping, with the peak association in both analyses located at 33,338,602 bp. To identify candidate genes, we examined annotated genes in the 31–35 Mb interval of Ra03 with homology to known flowering regulators. Ten genes were highlighted as candidates based on putative roles in flowering: Ruarg.3G323500 (UFC), Ruarg.3G329500 (BLH8), Ruarg.3G332300 (REV), Ruarg.3G334700 (CCCH-type zinc finger), Ruarg.3G337900 (SIZ1), Ruarg.3G338800 (AGL28), Ruarg.3G343800 (RAP2-7), Ruarg.3G346400 (UBP), Ruarg.3G349500 (AP2/B3-like), and Ruarg.3G349600 (VRN1) (Table 3).

In *Arabidopsis* and other model species, flowering is regulated by a network of transcriptional regulators and signaling pathways that integrate environmental and endogenous cues. Central “master regulators” such as *FLOWERING LOCUS T* (FT), *SUPPRESSOR OF OVEREXPRESSION OF CONSTANS 1* (SOC1), and *LEAFY* (LFY) coordinate the transition from vegetative to reproductive development, translating photoperiod, vernalization, and hormonal signals into activation of floral meristem identity genes (Song et al., 2013; Dinh et al., 2017). FT acts as a mobile florigen produced in leaves and transported to the shoot apical meristem (SAM), triggering flowering, while *TERMINAL FLOWER1* (TFL1) acts antagonistically to maintain vegetative growth (Koskela et al., 2012; Mimida et al., 2011).

Although a TFL1 homolog is present in blackberry (Ruarg.6G170000 on Ra06), it is located outside the PF locus, ruling it out as the causal gene.

Among the ten candidate genes in the locus, four harbor genetic variants that segregate with the PF phenotype (Table 4). Ruarg.3G329500, a homolog of the BEL1-like homeodomain 8 (BLH8, also called POUND-FOOLISH) plant homeobox transcription factor coding gene, contains a single intronic InDel (32,625,674 bp) distinguishing PF and FF genotypes. In Arabidopsis, BLH8, works together with another BELL1-like homeodomain protein, PENNYWISE (PNY), to maintain the shoot apical meristem (SAM) and regulate floral transition (Ung et al. 2011).

Another homeobox TF found in this region, Ruarg.3G332300, also harbors three intronic SNP variants and one InDel within 1 Kb upstream of the gene that segregate with PF. Ruarg.3G332300 is a homolog of the Class III Homeodomain-leucine zipper protein REVOLUTA (REV), which is necessary for shoot apical meristem development (Talbert et al. 1995) and required for the initiation of both lateral shoot meristems and flower meristems (Otsuga et al. 2001).

One homolog of a ubiquitin-specific protease, Ruarg.3G346400, was also found in the genomic region associated with the PF trait. UBP has been implicated in flowering time regulation through the photoperiod pathway. Mutations in UBP12 and UBP13 promote earlier expression of *CONSTANS*, which leads to increased expression of *FT* and earlier flowering in *Arabidopsis* (Cui et al. 2013). Ruarg.3G346400, contains two intronic variants, three SNP variants in the 3’UTR and two SNP variants within 1 Kb downstream of the gene differentiating PF and FF genotypes (Table 4). While these changes may have functional consequences, Ruarg.3G346400 is located over 1.2 Mb from the GWAS peak, making it a less likely causal gene. Moreover, UBP acts as a flowering promoter; a loss-of-function mutation would be expected to delay flowering rather than confer the recessive PF phenotype. This contrasts with the expected behavior of a mutation in a flowering repressor, consistent with the recessive inheritance of primocane fruiting.

By contrast, Ruarg.3G334700, a CCCH-type zinc finger gene, is positioned just 68 Kb from the GWAS peak and harbors the highest density of PF-associated variants within the locus: three intronic variants, two 3′ UTR variants, six variants within 1 kb downstream, and one synonymous exonic SNP. Members of the CCCH zinc finger family have been implicated in flowering regulation across multiple species. In *Arabidopsis*, KHZ1 and KHZ2 promote floral transition, with double mutants displaying delayed flowering and overexpression leading to early flowering (Yan et al. 2017). Similar flowering-promoting roles have been observed for AaZFP3 in *Adonis amurensis* and CpC3H3 in *Chimonanthus praecox* (Wang et al. 2022; Liu et al. 2022), whereas overexpression of the alfalfa MsZFN gene led to delayed flowering (Chao et al. 2014), highlighting that CCCH proteins can act as either promoters or repressors depending on context. Importantly, FF blackberries are short-day plants, while most functional data on these homologs come from long-day species, which may alter regulatory networks controlling flowering. The close proximity of Ruarg.3G334700 to the GWAS peak, coupled with the concentration of PF-associated variants, makes it an interesting candidate for the PF phenotype in blackberry.

The remaining six genes in the 31–35 Mb interval [Ruarg.3G323500 (UFC), Ruarg.3G337900 (SIZ1), Ruarg.3G338800 (AGL28), Ruarg.3G343800 (RAP2-7), Ruarg.3G349500 (AP2/B3-like), and Ruarg.3G349600 (VRN1)] have annotated roles consistent with flowering regulation. UFC is located upstream of FLC, a major flowering repressor in *Arabidopsis*, although UFC’s direct role in flowering remains unclear, and neither of the publicly available blackberry genomes harbors a clear FLC homolog (Finnegan et al. 2004; Aljaser et al. 2022). SIZ1 acts as a floral repressor via SUMOylation of flowering regulators (Jin et al. 2008; Miura et al. 2013; Son et al. 2014). AGL28 and RAP2-7 are involved in promoting floral meristem identity (Agarwal and Khurana 2019; Du et al. 2020; Li et al. 2025; Yoo et al. 2006; Zhang et al. 2015), and AP2/B3-like transcription factors can modulate the photoperiodic pathway, promoting or repressing flowering through direct binding to the promoters of SOC1 and FT (Hu et al. 2021; Yu et al. 2020). Despite their plausible functions, none of these genes harbor variants that segregate with the PF trait, suggesting they are unlikely to be the primary causal gene in this population.

Nonetheless, their presence underscores the complexity of the PF locus and highlights potential regulatory interactions that could modulate flowering in tetraploid blackberry.

### Diagnostic KASP Markers for PF

The KASP markers developed in this study, *PF1* and *PF2*, demonstrated strong predictive ability for the PF phenotype, correctly classifying 96.8% and 96.5% of genotypes, respectively, across a large and diverse validation panel. These are among the first diagnostic molecular markers developed for a major agronomic trait in blackberry. Previously, the only other diagnostic markers linked to a trait of economic importance were those associated with the prickle-free cane phenotype (Johns et al. 2025).

The development of reliable diagnostic markers for PF represents a major advance for blackberry breeding. Although PF cultivars offer significant production advantages, including extended harvest seasons, the ability to schedule fruiting through primocane management, avoidance of winter injury, and adaptation to low-chill environments, they generally lag behind elite FF cultivars in fruit quality, firmness, and flavor (Clark et al. 2008). Improving PF germplasm therefore requires crosses between high-quality FF selections and PF parents. Because PF is recessively inherited in a multisomic tetraploid background, only one in six seedlings (≈17%) from a duplex FF (CCTT) × PF (TTTT) cross is expected to express the PF phenotype. This low frequency greatly increases the population size, space, and time required to recover PF progeny with elite fruit quality. Reliable markers enable breeders to identify FF parents that carry PF alleles and to cull FF seedlings prior to planting, thereby enriching the proportion of PF individuals in segregating populations and accelerating the development of high-quality PF cultivars.

The PF1 and PF2 markers were broadly effective across germplasm from multiple breeding programs, correctly predicting the phenotype of 37 of 39 selections from Hortifrut Genetics Ltd., 73 of 76 selections from the USDA-ARS HCPGIR program, and all accessions from the USDA-ARS-NCGR collection. This demonstrates that the markers are robust and transferable beyond the UADA breeding program, a critical feature for adoption in both public and private breeding efforts.

Most inconsistencies between phenotypic and genotypic predictions were observed in population 1937. This is likely due to challenges in scoring PF among seedlings, which were planted at high density, where primocanes from neighboring plants can intermingle. Additionally, basal flowering can sometimes be mistaken for true primocane fruiting when scoring individual plants. Such phenotyping ambiguity likely explains a substantial portion of the observed discrepancies between marker prediction and field phenotype. Three UADA selections (S307, S441, and S443) were scored as PF but were predicted to carry at least one FF allele at both PF1 and PF2 loci. All three share the same FF parent (S34), which displays strong basal flowering on floricanes late in the season when primocanes are actively flowering. These half-sib selections consistently exhibited less than 30% of primocanes flowering in any given year, suggesting that their ambiguous phenotype may result from alleles at a secondary locus influencing basal or partial primocane flowering, rather than recombination at the primary PF locus.

Both *PF1* and *PF2* markers target SNPs within an intron of Ruarg.3G335600, a homolog of β-galactosidase 8 (BGAL8). As this gene has no known role in flowering, the associated variants are unlikely to be causal. This raises the possibility that a small number of true recombinants exist in the population, reflecting historical crossover events within the linkage disequilibrium block surrounding the PF locus. Overall, the *PF1* and *PF2* markers performed exceptionally well across diverse germplasm and provide a powerful, easy-to-use tool for accelerating selection in blackberry breeding programs. These markers are already being used to identify PF allele carriers among elite FF parents, plan crosses, and cull non-PF seedlings before planting in the UADA breeding program. Broader implementation of these diagnostic markers will enhance breeding efficiency across programs and facilitate the continued improvement of fruit quality and yield in primocane-fruiting blackberry cultivars.

## Conclusions

This study provides strong evidence that a single recessive locus between 31 and 35 Mb on chromosome Ra03 controls the PF trait in tetraploid blackberry. The identification of this locus through both GWAS and linkage mapping represents a major advance in understanding the genetic basis of PF, a trait of exceptional economic importance in blackberry and raspberry whose genetic control has remained largely unresolved. Within this region, several candidate genes with annotated roles in flowering regulation were identified, and allele mining using whole-genome resequencing data revealed numerous variants distinguishing PF and FF genotypes. Notably, variants in the 3′ untranslated region of a CCCH-type zinc finger gene, located ∼68 kb from the GWAS peak, emerged as a particularly promising candidate for further study. The availability of a high-quality, chromosome-scale genome annotation enabled detailed investigation of gene content and allelic variation within the PF locus, laying the groundwork for future functional analyses. Gene expression profiling of shoot apical meristems and targeted functional validation will be critical to determine causal variants for PF. Finally, the development of two highly accurate KASP markers (*PF1* and *PF2*) tightly linked to the PF locus provides practical tools for marker-assisted selection, enabling more efficient development of high-quality PF cultivars. Together, these results advance our understanding of flowering regulation in short-day Rosaceae species and provide key resources for both fundamental and applied blackberry genetics.

## Data availability

The updated genome annotation for *R. argutus* cv. ‘Hillquist’ v.1.2 is available on Phytozome (https://phytozome-next.jgi.doe.gov/info/Rargutus_v1_2) and can also be accessed at the Genome Database for Rosaceae (tfGDR1090; https://www.rosaceae.org/publication_datasets). Raw Isoseq and Illumina transcriptome data for *R. argutus* cv. Hillquist are available at NCBI under Bioproject IDs PRJNA830911and PRJEB36280 (BioSamples SAMEA6502409, SAMEA6502410, SAMEA6502411, SAMEA6502412, and SAMEA6502413), respectively. The whole-genome resequencing data used in this study are also publicly available at NCBI under Bioproject PRJNA1002337. The Capture-Seq derived SNP data used in GWAS are available in the Genome Database for Rosaceae repository (accession number tfGDR1069). All other data used in this study are presented in this manuscript and its supplementary files.

## Acknowledgments

The authors would like to thank Dr. Jackie Lee and the entire staff of the UADA Fruit Research Station for their work managing the plants used in this study. The bioinformatic analyses were supported by the Arkansas High Performance Computing Center, which is funded through multiple National Science Foundation grants and the Arkansas Economic Development Commission. Finally, we would like to thank LGC Genomics for their work on KASP primer design and the genotypic screening of the validation panel.

## Authors’ Contributions

AS collected phenotype data, performed GWAS and linkage analyses, analyzed WGS data, selected targets for KASP primer design, and drafted the manuscript in collaboration with MW. MW wrote the grant to support this research, supervised students and staff working on this project, and provided overall conceptual guidance. TB performed annotation analyses. TMC performed tetraploid SNP calling and bioinformatic analyses with CaptureSeq data. LN and CJ performed DNA extractions and quantification. IV conducted phenotyping, plant tissue collection, and genotyping in the biparental mapping population. JRC developed most of the advanced selections and created the biparental population used in the study. ET, MH, and NB contributed germplasm and phenotypic data for KASP marker validation. MH and NB also collaborated in Allegro® probe design and sequencing. MM developed MapPoly2 and collaborated with AS on linkage mapping. All authors contributed to the article and approved the submitted version.

## Supplementary Tables Legends

Supplementary Table 1. Blackberry selections and cultivars included in the genome-wide association analysis, their phenotypic description as Primocane-Fruiting (PF) or Floricane-Fruiting (FF) genotypes, and the allele dosage for the highest associated SNP markers at 33,338,602 and 33,338,650 bp on chromosome Ra03.

Supplementary Table 2. A total of 419 significant SNPs associated with the primocane-fruiting trait in the genome-wide association with -log10(p) values and effects for the simplex dominance (1-dom-ref) model.

Supplementary Table 3. Genotypic data for the biparental population ‘1937’ (113 progeny) derived from a cross between the FF genotype ‘S16’ and the PF genotype ‘S242’, along with their phenotypic classification as PF (Primocane-Fruiting) or FF (Floricane-Fruiting).

Supplementary Table 4. 53 SNPs highly linked with the Primocane-Fruiting phenotype in the blackberry biparental population’1937’. LOD score of the recombination fraction between each molecular marker and the PF phenotype was calculated, and a maximum LOD score was observed for the marker 3_33338602 .

Supplementary Table 5. List of genetic variants located within 1 Mb upstream and downstream of the GWAS peak associated with the primocane-fruiting trait on chromosome Ra03 differentiating Primocane-Fruiting (PF) and Floricane-Fruiting (FF) genotypes.

Supplementary Table 6. Allele dosage of 17 blackberry selections and cultivars, sequenced by whole-genome sequencing, for 12 SNP targets selected for KASP assay design to discriminate between PF and FF genotypes.

Supplementary Table 7. Primer sequence for Kompetitive Allele Specific Primer (KASP) assays targeting SNP distributed trough the genomic region associated with the PF trait on chromosome Ra03.

Supplementary Table 8. KASP markers validation panel. Genotypes used in the KASP validation study, phenotype for the Primocane-Fruiting (PF) trait (presence/absence), germplasm source, and information for the two most predictive KASP markers (PF1, Ra03:33338650 and PF2, Ra03:33,338,602), including FAM and HEX fluorescence value and allele dosage.

## Supplementary Figure Legends

**Supplementary Figure 1.**
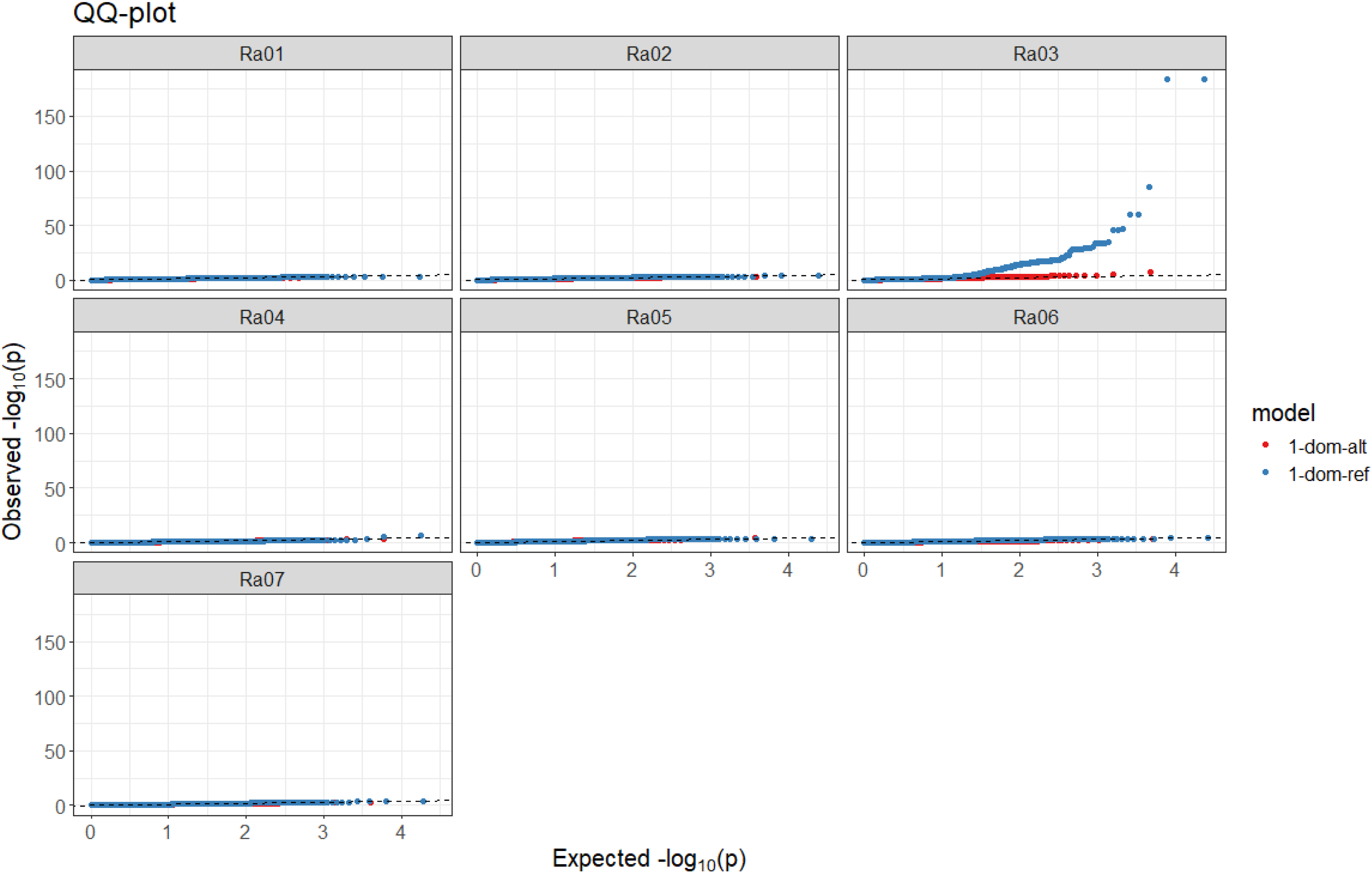
Chromosome-specific QQ-plots for the association analysis between the Primocane-Fruiting trait and SNPs under the simplex-dominant (1-dom) model.

**Supplementary Figure 2.**
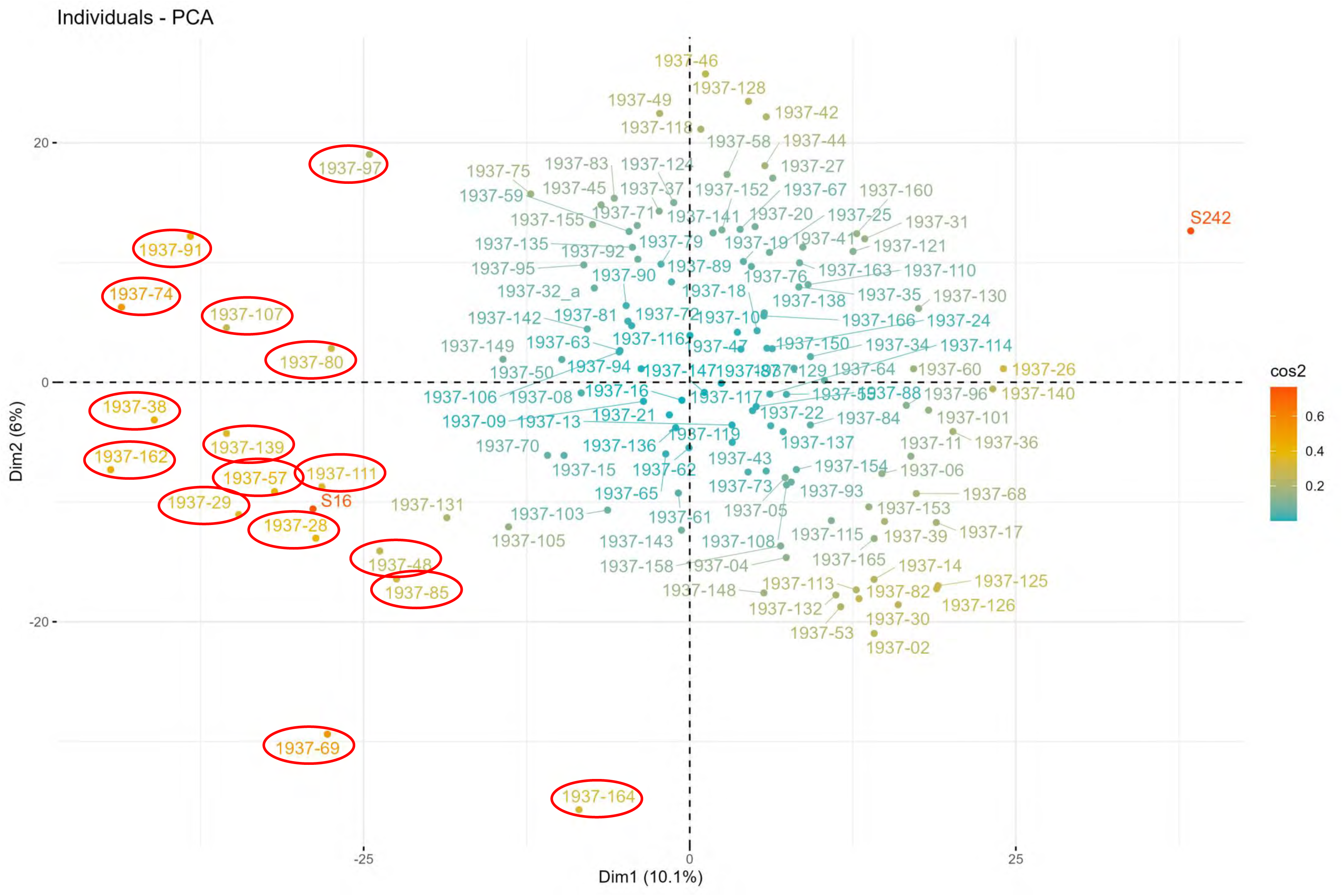
PCA for the biparental population ‘1937’ derived from the cross between the FF genotype ‘S16’ and the PF genotype ‘S242’. Samples highlighted in red were excluded from the genetic linkage analysis.

**Supplementary Figure 3.**
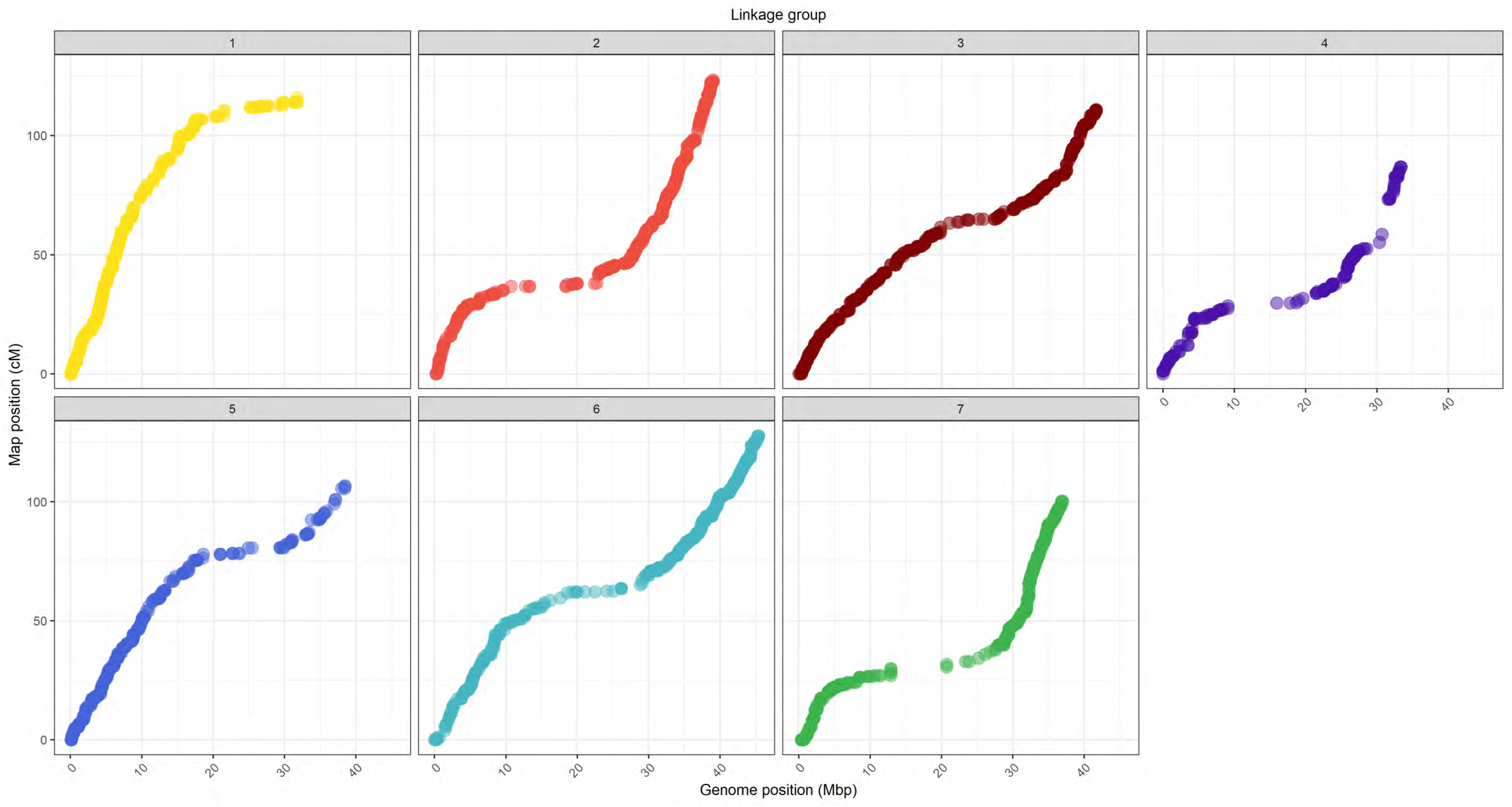
Comparison of the blackberry tetraploid genetic linkage map with the ‘Hillquist’ blackberry (*R. argutus*) physical map. As expected, seven linkage groups were identified for chromosomes Ra01 to Ra07. No noticeable inversions or translocations were detected.

**Supplementary Figure 4.**
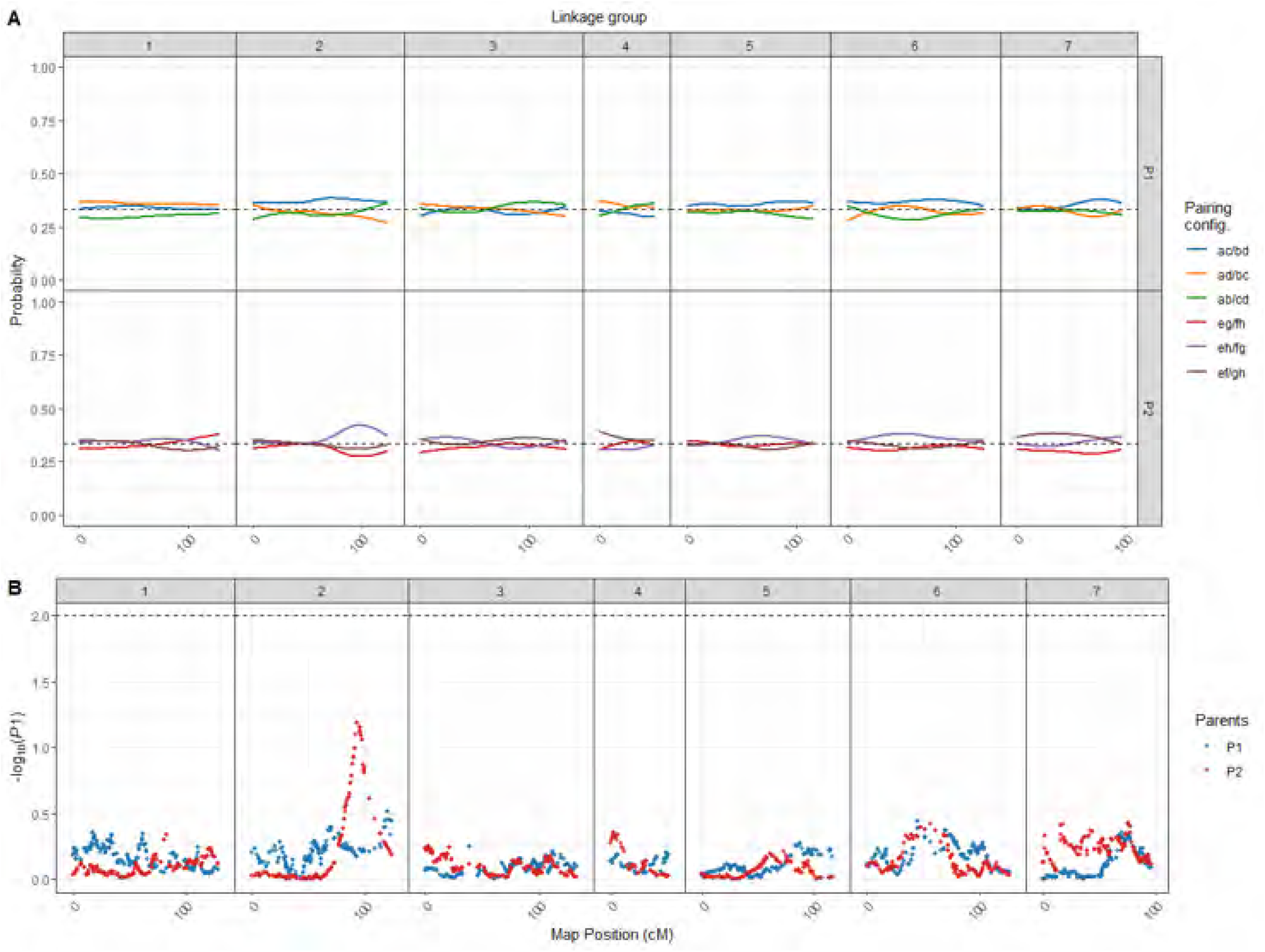
Preferential pairing profile in a tetraploid blackberry biparental population. (A) Probability profiles for homolog pairs in the parents ‘S16’ (P1) and ‘S242’ (P2) across 7 LGs. The dashed lines specify the pairing probability expected under random pairing. (B) *-log10(P)* of a χ 2 independence test for all possible homolog pairs where dashed lines indicate *P* < 10^−2^. No significant level of preferential pairing was observed across S16 or S242 parent.

## Literature Cited

Agarwal P, Khurana P. 2019. Overexpression of TaMADS from wheat promotes flowering by upregulating expression of floral promoters and provides protection against thermal stress. Plant Gene 17:100168. doi:10.1016/j.plgene.2018.100168.

Aljaser JA, Anderson NO, Noyszewski A. 2022. Discovery of UPSTREAM OF FLOWERING LOCUS C (UFC) and FLOWERING LOCUS C EXPRESSOR (FLX) in Gladiolus ×hybridus, G. dalenii. Ornamental Plant Research 2:13 doi: 10.48130/OPR-2022-0013

Bao W, Kojima KK, Kohany O. 2015. Repbase Update, a database of repetitive elements in eukaryotic genomes. Mobile DNA. 6:11. doi: 10.1186/s13100-015-0041-9

Bolger AM, Lohse M, Usadel B. 2014. Trimmomatic: a flexible trimmer for Illumina sequence data. Bioinformatics. 30(15):2114–2120. doi:10.1093/bioinformatics/btu170.

Bourke PM, Voorrips RE, Visser RGF, Maliepaard C. 2015. The double-reduction landscape in tetraploid potato as revealed by a high-density linkage map. Genetics 201(3):853–863. doi:10.1534/genetics.115.181008

Brůna T, Aryal R, Dudchenko O, Sargent DJ, Mead D, Buti M, et al. 2022. A chromosome-length genome assembly and annotation of blackberry (Rubus argutus, cv. ‘Hillquist’). G3 Genes|Genomes|Genetics. 13:jkac289. doi:10.1093/g3journal/jkac289.

Cappai F, Amadeu RR, Benevenuto J, Cullen R, Garcia A, Grossman A, et al. 2020. High-Resolution Linkage Map and QTL Analyses of Fruit Firmness in Autotetraploid Blueberry. Front Plant Sci. 11:562171. doi:10.3389/fpls.2020.562171.

Castro P, Stafne ET, Clark JR, Lewers KS. 2013. Genetic map of the primocane-fruiting and thornless traits of tetraploid blackberry. Theoretical and Applied Genetics. 126(10):2521–2532. doi:10.1007/s00122-013-2152-3.

Chae L, Kim T, Nilo-Poyanco R, Rhee SY. 2014. Genomic signatures of specialized metabolism in plants. Science. 344(6183):510–513. doi: 10.1126/science.1252076.

Chao Y, Zhang T, Yang Q, Kang J, Sun Y, Gruber MY, Qin Z, 2014. Expression of the alfalfa CCCH-type zinc finger protein gene MsZFN delays flowering time in transgenic Arabidopsis thaliana. Plant Sci 215–216:92–99. doi:10.1016/j.plantsci.2013.10.012

Chizk TM, Clark JR, Johns C, Nelson L, Ashrafi H, Aryal R, Worthington ML. 2023. Genome-wide association identifies key loci controlling blackberry postharvest quality. Front. Plant Sci. 14:1182790. doi: 10.3389/fpls.2023.1182790

Clark JR, Stafne ET, Hall HK, Finn CE, 2007. Blackberry breeding and genetics. In: Janick J (ed) Plant Breeding Reviews, 29th edn. pp 19–144.

Clark JR, 2008. Primocane-fruiting Blackberry Breeding. HortScience 43(6):1637–1639. doi:10.21273/HORTSCI.43.6.1637

Clark, JR, Finn CE. 2014. Blackberry cultivation in the world. Rev. Bras. Frutic. 36(1): 46–57. doi:10.1590/0100-2945-445/13

Clark JR, Moore JN, Lopez-Medina J, Finn C, Perkins-Veazie P. 2005. ’Prime-Jan’ (’APF-8’) and’Prime-Jim’ (’APF-12’) primocane-fruiting blackberries. HortScience. 40(3):852–855. doi:10.21273/HORTSCI.40.3.852.

Cui X, Lu F, Li Y, Xue Y, Kang Y, Zhang S, Qiu Q, et al. 2013. Ubiquitin-specific proteases UBP12 and UBP13 act in circadian clock and photoperiodic flowering regulation in Arabidopsis. Plant Physiol. 162(2):897–906. doi:10.1104/pp.112.213009.

Danecek P, Auton A, Abecasis G, Albers C, Banks E, DePristo MA, Handsaker RE, Lunter G, Marth GT, Sherry ST, McVean G, Durbin R, 1000 Genomes Project Analysis Group. 2011. The variant call format and VCFtools. Bioinformatics. 27(15):2156–2158. doi:10.1093/bioinformatics/btr330.

Dinh JL, Farcot E, Hodgman C, 2017. The logic of the floral transition: reverse-engineering the switch controlling the identity of lateral organs. PLoS Comput Biol. 13(9): e1005744. doi:10.1371/journal.pcbi.1005744

Du SS, Li L, Li L, Wei X, Xu F, Xu P, Wang W, Xu P, et al. 2020. Photoexcited Cryptochrome2 Interacts Directly with TOE1 and TOE2 in Flowering Regulation. Plant Physiol 184(1):487–505. doi:10.1104/pp.20.00486

Duitama J, Quintero JC, Cruz DF, Quintero C, Hubmann G, Foulquie-Moreno MR, Verstrepen KJ, Thevelein JM, Tohme J. 2014. An integrated framework for discovery and genotyping of genomic variants from high-throughput sequencing experiments. Nucleic Acids Res. 42(6):e44. doi:10.1093/nar/gkt1381.

Ferrão LFV, Benevenuto J, Oliveira IB, de Bem Oliveira L, Cellon C, Olmstead J, et al. 2018. Insights into the genetic basis of blueberry fruit-related traits using diploid and polyploid models in a GWAS context. Front. Ecol. Evol. 6:107. doi:10.3389/fevo.2018.00107

Fernandez G, McWhirt A, Bradish C. 2023. Southeast Regional Caneberry Production Guide. NC State Extension Publications. https://content.ces.ncsu.edu/southeast-regional-caneberry-production-guide (Accessed October 28, 2025).

Finnegan EJ, Sheldon CC, Jardinaud F, Peacock WJ, Dennis ES. 2004. A cluster of Arabidopsis genes with a coordinate response to an environmental stimulus. Curr Biol 14:911–916. doi:10.1016/j.cub.2004.04.045

Flynn JM, Hubley R, Goubert C, Rosen J, Clark AG, Feschotte C, Smit AF. 2020. RepeatModeler2 for automated genomic discovery of transposable element families. Proc Natl Acad Sci U S A. 117(17):9451–9457. doi: 10.1073/pnas.1921046117.

Garrison E, Marth G. 2012.Haplotype-based variant detection from short-read sequencing. arXiv:1207.3907. doi: 10.48550/arXiv.1207.3907, preprint: not peer reviewed.

Gerard D. 2021. Pairwise linkage disequilibrium estimation for polyploids. Mol Ecol Resour. 21(4):1230–1242. doi:10.1111/1755-0998.13349.

Gerard D, Ferrão LFV, Garcia AAF, Stephens M. 2018. Genotyping polyploids from messy sequencing data. Genetics. 210:789–807. doi:10.1534/genetics.118.301468.

Godwin C, Chizk TM, Johns C, Nelson L, Threlfall R, Clark JR, Worthington ML, 2025. Genetic control of sweetness and acidity in blackberry. Front. Plant Sci. 16. doi:10.3389/fpls.2025.1569492.

Haas BJ, Delcher AL, Mount SM, Wortman JR, Smith RK Jr, Hannick LI, Maiti R, Ronning CM, Rusch DB, Town CD, Salzberg SL, White O. 2003. Improving the Arabidopsis genome annotation using maximal transcript alignment assemblies. Nucleic Acids Res. 31(19):5654–5666. doi:10.1093/nar/gkg770.

Hu H, Tian S, Xie G, Liu R, Wang N, Li S, He Y, Du J. 2021. TEM1 combinatorially binds to FLOWERING LOCUS T and recruits a Polycomb factor to repress the floral transition in Arabidopsis. PNAS 118(35):e2103895118. doi: 10.1073/pnas.2103895118.

Hubley R, Finn RD, Clements J, Eddy SR, Jones TA, Bao W, Smit AF, Wheeler TJ. 2016. The Dfam database of repetitive DNA families. Nucleic Acids Res. 44(D1):D81–9. doi: 10.1093/nar/gkv1272.

Jibran R, Spencer J, Fernandez G, Monfort A, Mnejja M, Dzierzon H, Tahir J, Davies K, Chagné D, Foster TM. 2019. Two Loci, RiAF3 and RiAF4, Contribute to the Annual-Fruiting Trait in Rubus. Front. Plant Sci. 10:1341. doi: 10.3389/fpls.2019.01341.

Jin JB, Jin YH, Lee J, Miura K, Yoo CY, et al. 2008. The SUMO E3 ligase, AtSIZ1, regulates flowering by controlling a salicylic acid-mediated floral promotion pathway and through affects on FLC chromatin structure. Plant J 53:530–540. doi:10.1111/j.1365-313X.2007.03359.x.

Johns CA, Silva A, Chizk TM, Nelson L, Clark JR, Aryal R, Ashrafi H, Thompson E, Hardigan M, Worthington ML. 2025. Genetic control of prickles in tetraploid blackberry. G3 GENES|GENOMES|GENETICS. 15(6):jkaf065. doi:10.1093/g3journal/jkaf065.

Jones P, Binns D, Chang HY, Fraser M, Li W, McAnulla C, McWilliam H, Maslen J, Mitchell A, Nuka G, et al. 2014. InterProScan 5: genome-scale protein function classification. Bioinformatics. 30(9):1236–40. doi: 10.1093/bioinformatics/btu031.

Karp PD, Latendresse M, Caspi R. 2011. The pathway tools pathway prediction algorithm. Stand Genomic Sci. 5(3):424–429. doi: 10.4056/sigs.1794338.

Koskela EA, Mouhu K, Albani MC, Kurokura T, Rantanen M, Sargent DJ, Battey NH, Coupland G, Elomaa P, Hytönen T, 2012. Mutation in TERMINAL FLOWER1 reverses the photoperiodic requirement for flowering in the wild strawberry Fragaria vesca. Plant Physiol 159:1043–1054. doi:10.1104/pp.112.196659.

Lau J, Young EL, Collins S, Windham MT, Klein PE, Byrne DH, Riera-Lizarazu O. 2022. Rose Rosette Disease Resistance Loci Detected in Two Interconnected Tetraploid Garden Rose Populations. Front. Plant Sci. 13:916231. doi:10.3389/fpls.2022.916231.

Lee W-P, Stromberg MP, Ward A, Stewart C, Garrison EP, Marth GT. 2014. MOSAIK: a hash-based algorithm for accurate next-generation sequencing short-read mapping. PLoS One. 9:e90581. doi:10.1371/journal.pone.0090581.

Li H, Durbin R. 2009. Fast and accurate short read alignment with Burrows–Wheeler transform. Bioinformatics. 25(14):1754–1760. doi:10.1093/bioinformatics/btp324.

Li Y, Yang H, Jia P, Li Z, Wang Y, Jiang Y, He X, Wen B, Huo C, Zhang W, Chai W, Yan S, Zhang J, 2025. The MADS-Box Transcription Factor BoAGL8 is Involved in Positive Regulation of Flowering in Broccoli. SSRN. doi:10.2139/ssrn.5371165.

Liu H, Xiao S, Sui S, et al., 2022. A tandem CCCH type zinc finger protein gene CpC3H3 from Chimonanthus praecox promotes flowering and enhances drought tolerance in Arabidopsis. BMC Plant Biol 22:506. doi:10.1186/s12870-022-03877-2.

Lopez-Medina J, Moore JN, McNew RW. 2000. A proposed model for inheritance of primocane fruiting in tetraploid erect blackberry. Journal of the American Society for Horticultural Science. 125(2):217–221. doi:10.21273/jashs.125.2.217.

Oloka BM, da Silva Pereira G, Amankwaah VA, Mollinari M, Pecota KV, Yada B, et al. 2021. Discovery of a major QTL for root-knot nematode (Meloidogyne incognita) resistance in cultivated sweetpotato (Ipomoea batatas). Theor Appl Genet. 134:1945–1955. 10.1007/s00122-021-03797-z.

Otsuga D, DeGuzman B, Prigge MJ, Drews GN, Clark SE. 2001. REVOLUTA regulates meristem initiation at lateral positions. Plant J 25:223–236. doi:10.1046/j.1365-313x.2001.00959.x.

Paudel D, Parrish SB, Peng Z, Parajuli S, Deng Z, 2025. A chromosome-scale and haplotype-resolved genome assembly of tetraploid blackberry (Rubus L. subgenus Rubus Watson). Hortic. Res. 12(6):uhaf052. doi:10.1093/hr/uhaf052.

Porebski S, Bailey LG, Baum BR. 1997. Modification of a CTAB DNA extraction protocol for plants containing high polysaccharide and polyphenol components. Plant Mol Biol Report. 15(1):8–15. doi:10.1007/bf02772108.

Poplin R, Ruano-Rubio V, DePristo MA, Fennell TJ, Carneiro MO, Van der Auwera GA, Kling DE, Gauthier LD, Levy-Moonshine A, Roazen D, et al. 2018. Scaling accurate genetic variant discovery to tens of thousands of samples. bioRxiv. doi:10.1101/201178.

Rosyara UR, De Jong WS, Douches DS, Endelman JB. 2016. Software for genome-wide association studies in autopolyploids and its application to potato. Plant Genome. 9(2):plantgenome2015.2008.0073. doi:10.3835/plantgenome2015.08.0073.

Mather K.1936. Segregation and linkage in autotetraploids. Journal of Genetics. 32, 287–314. doi:10.1007/BF02982683

Mimida N, Ureshino A, Tanaka N, Ito A, Ichikawa H, Souda K, et al. 2011. Expression patterns of several floral genes during flower initiation in the apical buds of apple (Malus × domestica Borkh.) revealed by in situ hybridization. Plant Cell Rep. 30(9):1485–1492. doi:10.1007/s00299-011-1057-3.

Mistry J, Chuguransky S, Williams L, Qureshi M, Salazar GA, Sonnhammer ELL, Tosatto SCE, Paladin L, Raj S, Richardson LJ, et al. 2021. Pfam: The protein families database in 2021. Nucleic Acids Res. 49(D1):D412–D419. doi: 10.1093/nar/gkaa913.

Miura K, Okamoto H, Okuma E, Shiba H, Kamada H, Hasegawa PM, Murata Y, 2013. SIZ1 deficiency causes reduced stomatal aperture and enhanced drought tolerance via controlling salicylic acid-induced accumulation of reactive oxygen species in Arabidopsis. Plant J 73: 91–104. doi:10.1111/tpj.12014.

Mollinari M, Garcia AA. 2019. Linkage analysis and haplotype phasing in experimental autopolyploid populations with high ploidy level using Hidden Markov models. G3 Genes|Genomes|Genetics. 9(10): 3297–3314. doi:10.1534/g3.119.400378

Mollinari M, Olukolu BA, da Silva Pereira G, Khan A, Gemenet D, Yencho GC, Zeng Z-B. 2020. Unraveling the Hexaploid Sweetpotato Inheritance Using Ultra-Dense Multilocus Mapping. G3 Genes|Genomes|Genetics. 10(1):281–292. doi:10.1534/g3.119.400620

Saavedra-Díaz C, Trujillo-Montenegro JH, Jaimes HA, Londoño A, et al. 2024. Genetic association analysis in sugarcane (Saccharum spp.) for sucrose accumulation in humid environments in Colombia. BMC Plant Biol.24(1):570. doi:10.1186/s12870-024-05233-y

Salamov AA, Solovyev VV. 2000. Ab initio gene finding in Drosophila genomic DNA. Genome Res. 10(4):516–522. doi: 10.1101/gr.10.4.516.

Schläpfer P, Zhang P, Wang C, Kim T, Banf M, Chae L, Dreher K, Chavali AK, Nilo-Poyanco R, Bernard T, et al. 2017. Genome-wide prediction of metabolic enzymes, pathways, and gene clusters in plants. Plant Physiol. 173(4):2041–2059. doi: 10.1104/pp.16.01942.

Sharma SK, MacKenzie K, McLean K, Dale F, Daniels S, Bryan GJ. 2018. Linkage Disequilibrium and Evaluation of Genome-Wide Association Mapping Models in Tetraploid Potato. G3 (Bethesda) 8(10):3185-3202. doi:10.1534/g3.118.200377.

da Silva Pereira G, Mollinari M, Schumann MJ, et al., 2021. The recombination landscape and multiple QTL mapping in a Solanum tuberosum cv. ‘Atlantic’-derived F1 population. Heredity 126:817–830. doi:10.1038/s41437-021-00416-x.

da Silva Pereira G, Mollinari M, Qu X, Thill C, Zeng Z-B, Haynes K, Yencho GC. 2021. Quantitative Trait Locus Mapping for Common Scab Resistance in a Tetraploid Potato Full-Sib Population. Plant Dis. 105(10):3048–3054. doi:10.1094/PDIS-10-20-2270-RE

Slater GSC, Birney E. 2005. Automated generation of heuristics for biological sequence comparison. BMC Bioinformatics. 6:31. doi: 10.1186/1471-2105-6-31.

Smit A, Hubley R, Green P. RepeatMasker Open-4.0; 2013. https://www.repeatmasker.org.

Sommer M J, Zimin A V, Salzberg S L. 2025. PSAURON: a tool for assessing protein annotation across a broad range of species. NAR Genomics and Bioinformatics 7(1): lqae189. doi:10.1093/nargab/lqae189.

Son GH, Park BS, Song JT, Seo HS, 2014. FLC-mediated flowering repression is positively regulated by sumoylation. J Exp Bot 65:339–351. doi:10.1093/jxb/ert383.

Song GQ, Walworth A, Zhao D, Jiang N, Hancock JF. 2013. The Vaccinium corymbosum FLOWERING LOCUS T-like gene (VcFT): a flowering activator reverses photoperiodic and chilling requirements in blueberry. Plant Cell Rep. 32(11):1759–1769. doi: 10.1007/s00299-013-1489-z

Stanke M, Schöffmann O, Morgenstern B, Waack S. 2006. Gene prediction in eukaryotes with a generalized hidden Markov model that uses hints from external sources. BMC Bioinformatics. 7:62. doi: 10.1186/1471-2105-7-62.

Talbert PB, Adler HT, Parks DW, Comai L. 1995. The REVOLUTA gene is necessary for apical meristem development and for limiting cell divisions in the leaves and stems of Arabidopsis thaliana. Development 121:2723–2735. doi:10.1242/dev.121.9.2723.

Ung N, Lal S, Smith HM. 2011. The role of PENNYWISE and POUND-FOOLISH in the maintenance of the shoot apical meristem in Arabidopsis. Plant Physiol 156:605–614. doi:10.1104/pp.110.171462.

VanBuren R, Bryant D, Bushakra JM, Vining KJ, Edger PP, Rowley ER, Priest HD, Michael TP, Lyons E, Filichkin SA, et al. 2016. The genome of black raspberry (Rubus occidentalis). Plant J. 87:535–547. doi:10.1111/tpj.13215.

Vasimuddin M, Misra S, Li H, Aluru S. 2019. Efficient architecture-aware acceleration of BWA-mem for multicore systems. 2019. IEEE International Parallel and Distributed Processing Symposium (IPDPS). 314–324. doi:10.1109/IPDPS.2019.00041

Vaughn I, Silva A, Johns C, Nelson L, Worthington M. 2023. Validation of a Diagnostic Marker for Primocane-Fruiting in Blackberry. Discovery, The Student Journal of Dale Bumpers College of Agricultural, Food and Life Sciences, 24(1). Retrieved from https://scholarworks.uark.edu/discoverymag/vol24/iss1/13.

VanBuren R, Bryant D, Bushakra JM, Vining KJ, Edger PP, Rowley ER, et al. 2016. The genome of black raspberry (Rubus occidentalis). Plant J. 87(6):535–547. doi:10.1111/tpj.13215.

Wang M, Zhang H, Dai S, Feng S, Gong S, Wang J, Zhou A. 2022. AaZFP3, a novel CCCH-type zinc finger protein from Adonis amurensis, promotes early flowering in Arabidopsis by regulating the expression of flowering-related genes. Int J Mol Sci 23:8166. doi:10.3390/ijms23158166.

Wilson S, Zheng C, Maliepaard C, Mulder HA, Visser RGF, van der Burgt A, van Eeuwijk F, 2021. Understanding the Effectiveness of Genomic Prediction in Tetraploid Potato. Front. Plant Sci. 12:672417. doi:10.3389/fpls.2021.672417.

Wickham, H. 2016. ggplot2: Elegant Graphics for Data Analysis. Springer-Verlag New York. ISBN 978-3-319-24277-4

Wu TD, Nacu S. 2010. Fast and SNP-tolerant detection of complex variants and splicing in short reads. Bioinformatics. 26(7):873–881. doi: 10.1093/bioinformatics/btq057.

Yan Z, Jia J, Yan X, Shi H, Han Y. 2017. Arabidopsis KHZ1 and KHZ2, two novel non-tandem CCCH zinc-finger and K-homolog domain proteins, have redundant roles in the regulation of flowering and senescence. Plant Mol Biol 95:549–565. doi:10.1007/s11103-017-0667-8.

Yoo SK, Lee JS, Ahn JH, 2006. Overexpression of AGAMOUS-LIKE 28 (AGL28) promotes flowering by upregulating expression of floral promoters within the autonomous pathway. Biochem Biophys Res Commun 348:929–936. doi:10.1016/j.bbrc.2006.07.121

Zhang B, Wang L, Zeng L, Zhang C, Ma H. 2015. Arabidopsis TOE proteins convey a photoperiodic signal to antagonize CONSTANS and regulate flowering time. Genes Dev 29(9):975–987. doi:10.1101/gad.251520.114.

